# Phenome-wide Investigation of Health Outcomes Associated with Genetic Predisposition to Loneliness

**DOI:** 10.1101/468835

**Authors:** Abdel Abdellaoui, Sandra Sanchez-Roige, Julia Sealock, Jorien L. Treur, Jessica Dennis, Pierre Fontanillas, Sarah Elson, The 23andme Research Team, Michel Nivard, Hill Fung Ip, Matthijs van der Zee, Bart Baselmans, Jouke Jan Hottenga, Gonneke Willemsen, Miriam Mosing, Li Yu, Nancy L. Pedersen, Najaf Amin, Cornelia M van Duijn, Ingrid Szilagyi, Henning Tiemeier, Alexander Neumann, Karin Verweij, Stephanie Cacioppo, John T. Cacioppo, Lea K. Davis, Abraham A. Palmer, Dorret I. Boomsma

**Affiliations:** Department of Biological Psychology, Vrije Universiteit, Amsterdam, Netherlands; Department Psychiatry, Amsterdam UMC, University of Amsterdam, Amsterdam, the Netherlands; Department of Psychiatry, University of California San Diego, La Jolla, CA, USA; Vanderbilt Genetics Institute, Division of Genetic Medicine, Department of Medicine, Vanderbilt University, Nashville, TN, USA; School of Experimental Psychology, University of Bristol, Bristol, UK; MRC Integrative Epidemiology Unit, University of Bristol, Bristol, UK; 23andMe, Inc., Mountain View, CA, USA; Department of Neuroscience, Karolinska Institutet, Stockholm, Sweden; Department of Medical Epidemiology and Biostatistics, Karolinska Institutet, Stockholm, Sweden; Genetic Epidemiology Unit, Department of Epidemiology, Erasmus Medical Center, Rotterdam, Netherlands; Translational Epidemiology, Faculty Science, Leiden University, Leiden, the Netherlands; Department of Epidemiology, Erasmus Medical Center, Rotterdam, Netherlands; Department of Psychiatry, Erasmus Medical Center, Rotterdam, Netherlands; Department of Child and Adolescent Psychiatry/Psychology, Erasmus Medical Center, Rotterdam, Netherlands; Center for Cognitive and Social Neuroscience, Department of Psychology, The University of Chicago, Chicago, Illinois, USA

## Abstract

Humans are social animals that experience intense suffering when they perceive a lack of social connection. Modern societies are experiencing an epidemic of loneliness. While the experience of loneliness is universally human, some people report experiencing greater loneliness than others. Loneliness is more strongly associated with mortality than obesity, emphasizing the need to understand the nature of the relationship between loneliness and health. While it is intuitive that circumstantial factors such as marital status and age influence loneliness, there is also compelling evidence of a genetic predisposition towards loneliness. To better understand the genetic architecture of loneliness and its relationship with associated outcomes, we conducted a genome-wide association (GWAS) meta-analysis of loneliness (N=475,661), report 12 associated loci (two novel) and significant genetic correlations with 34 other complex traits. The polygenic basis for loneliness was significantly enriched for evolutionary constrained genes and genes expressed in specific brain tissues: (frontal) cortex, cerebellum, anterior cingulate cortex, and substantia nigra. We built polygenic scores based on this GWAS meta-analysis to explore the genetic association between loneliness and health outcomes in an independent sample of 18,498 individuals for whom electronic health records were available. A genetic predisposition towards loneliness predicted cardiovascular, psychiatric, and metabolic disorders, and triglycerides and high-density lipoproteins. Mendelian randomization analyses showed evidence of a causal, increasing, effect of body fat on loneliness, and a similar weaker causal effect of BMI. Our results provide a framework for ongoing studies of the genetic basis of loneliness and its role in mental and physical health.

## Introduction

Loneliness is a universal human experience that has been documented across cultures and generations. According to the evolutionary theory of loneliness^1^, this familiar painful feeling corresponds to an aversive response to a discrepancy between a people’s desired and perceived level of social connectedness.^2,3^ This definition, which emphasizes the *desired* level of social connection, highlights the difference between loneliness and solitude. Unlike solitude, the signal associated with loneliness has likely evolved to motivate humans and other social animals to seek and improve the salutary social connections needed to help them survive and reproduce.^4^ Loneliness serves as an emotional warning or signal that there is an emotional imbalance in one’s social network, regardless of the size of that network. Feeling lonely is also very common; about 5-30% of adults in Western populations report some degree of loneliness, while the actual prevalence may be higher since loneliness is stigmatized in many cultures.^5-7^

Multiple factors influence the individual differences in the experience of chronic loneliness.^1^ Most studies have focused on circumstantial factors such as marital status, age, and sex.^8-11^ However, there are also innate individual differences in the propensity to feel lonely. Heritability estimates based on twin and family data suggest that ~37% of the variation in loneliness levels is explained by genetic factors^12^. Studies using molecular genetic data provided evidence that the aggregate of common genetic variants account for 4-27% in individual differences in loneliness,^13-15^ and a recent genome-wide association study (GWAS) of social interaction and isolation in the UK Biobank sample has identified 15 common genetic variants associated with loneliness.^15^

Both social isolation and chronic high levels of loneliness are strongly correlated with negative health outcomes; chronic loneliness has a stronger association with early mortality than obesity.^16^ A long-running longitudinal study on physical and mental health, the Harvard Study of Adult Development, has concluded that the warmth of one’s relationships has the greatest impact on wellbeing and life satisfaction.^17^ Findings like these suggest that loneliness is a public health concern. While these studies demonstrate a clear and strong correlation between loneliness and increased morbidity and mortality, the causality and etiology of the relationship between loneliness and mental and physical health is unclear. For example, one possibility is that loneliness may cause poor health, or, alternatively, poor health may cause loneliness directly or indirectly, for example, by disrupting social networks.

Here, we first conducted a GWAS meta-analysis for loneliness on nearly half-a-million subjects of European descent from various cultural backgrounds, including the UK biobank (UKB), 23andMe (USA), the Health and Retirement Study (USA), the Netherlands Twin Register (NTR), and the Swedish Twin Registry (STR). We performed secondary analyses on the summary statistics^18^ using gene-based and LD score regression approaches to further elucidate the biological basis underlying the propensity to feel lonely and the genetic overlap between loneliness and complex human traits related to personality, cognition, reproduction, substance use, social connections, and physical and mental health. Next, we carried out a phenome wide association study (PheWAS). PheWAS have emerged as a method to screen for associations between genetic measures and a range of phenotypes, such as those measured in electronic health records (EHR).^19,20^ For certain phenotypes, EHR may provide more objective measures of physical and mental health than self-reported health data, which may not be readily known by patients (e.g., lab values) or can be distorted by mood and recall bias. Since the time of their original publications, the PheWAS approach has expanded beyond analysis of a single SNP to also include analysis of polygenic risk predictors.^21^ In this study, we constructed a polygenic predictor of loneliness using estimated SNP effects from the GWAS meta-analysis and performed a PheWAS on this polygenic score in the Vanderbilt University Medical Center (VUMC) EHR and associated biobank. Subsequent to this analysis, we analyzed a subset of quantitative traits that were significantly associated with the loneliness polygenic score in our PheWAS and known to be biomarkers for diagnoses. The goal of this analysis was to determine whether polygenic scores for loneliness were associated with known causal biomarkers. However, these analyses, like others that rely on genetic correlations, do not distinguish causal effects from pleiotropic effects. Therefore, we further tested for bidirectional causal relationships between loneliness and a selection of genetically correlated phenotypes using Mendelian randomization. We performed a comprehensive characterization of the polygenic contribution to the universal human experience of loneliness and extended this understanding to elucidate the genetic relationships between loneliness and health.

## Subjects and Methods

### Subjects & Phenotype

A total of 475,661 adult subjects from 7 different cohorts were included in the GWAS meta-analysis. An overview of subjects and phenotyping across cohorts can be found in Supplementary Table 1. The UK Biobank (UKB) dataset was the largest. UKB was the only cohort with a dichotomous phenotype (N_total_ = 413,337: 74,142 lonely and 339,195 non-lonely individuals). The other six cohorts had three types of continuous measures for loneliness: the sum of 9 items on a 4-point scale, the sum of 3 items on a 3-point scale, and 1 item on a 4-point scale.

### Genotyping and QC

Information on genotyping, imputation, and QC is given in Supplementary Table 2. In all cohorts, SNP data were imputed to either 1000 Genomes or the Haplotype Reference Consortium (HRC). SNPs remaining after QC ranged from 5.7 million to 14.1 million. Based on ancestry information derived from SNP data, only subjects with European descent were included.

### GWASs & Meta-Analysis

GWASs were performed in all seven cohorts separately, with the variables age, sex, family relationships, and ancestry-informative PCs as fixed effects (see Supplementary Table 2 for details). The UK Biobank dataset was split into three groups of unrelated individuals on which three separate categorical GWASs were run, and there were six continuous GWASs for the other six cohorts. The categorical GWASs (logistic regressions) on UKB were run on the following three groups: 1) the largest group of unrelated individuals with British ancestry (N_total_ = 332,991: 58,960 cases & 274,031 controls), 2) individuals with British ancestry that consist of family members of the first group (N_total_ = 57,865: 10,430 cases & 47,435 controls), and 3) individuals of Non-British European descent (N_total_ = 22,481: 4,752 cases & 17,729 controls). The two groups of GWASs (i.e., three categorical GWASs and six continuous GWASs) were first meta-analyzed separately using the multivariate approach described in Baselmans et al (2017)^22^. This approach controls for bias due to relatedness or sample overlap between GWASs by incorporating the cross-trait LD-score intercept (a measure for sample overlap) from LD-score regression (LDSC)^23^ as weights, which was especially necessary for the UK Biobank datasets, since the second dataset of unrelated individuals consisted of family members of the first dataset of unrelated individuals. The two meta-analyses (categorical and continuous) were then meta-analyzed using sample size-based weights to accounts for the respective differences of heritabilities, genetic correlation, and measurement scales of the categorical and continuous GWASs (see Demontis et al, 2017, for more details).^24^

### Follow-up Analyses

*Gene-based tests &* g*ene enrichment tests*: GWAS meta-analysis summary statistics were used to compute gene-based *p*-values in MAGMA^25^ for 18,125 protein coding genes using FUMA.^26^ MAGMA in FUMA was further used to test whether the effects of genes on loneliness were correlated with higher or lower gene-expression in a given tissue based on GTEx RNA-seq data.^26^ This was tested for 30 general tissue types and 53 more specific tissue types.

*LD-Score Regression Heritability Partitioning*: Stratified LD-score regression was carried out using LDSC in order to partition the heritability signal into specific cell-type groups or genomic annotations.^27,28^ This method requires the GWAS meta-analysis summary statistics, and LD information based on an external reference panel, for which we used the European populations from the HapMap 3 reference panel.

*S-PrediXcan*: S-PrediXcan^29^ uses reference panels with both measured gene expression and genotype data collected on the same individuals to build predictive models of gene expression in samples in which only genotype information is available. Predicted expression of genes for cases and controls can then be associated with phenotypic differences, yielding a gene-based test of association that incorporates transcriptional information. We used S-PrediXcan^29^ to predict gene expression levels in 10 brain tissues, and to test whether the predicted gene expression correlates with loneliness. Pre-computed tissue weights were employed from the Genotype-Tissue Expression (GTEx v7) project database (https://www.gtexportal.org/)^30^ as the reference transcriptome dataset. As input data, we included the loneliness GWAS meta-analysis summary statistics, transcriptome tissue data, and covariance matrices of the SNPs within each gene model (based on HapMap SNP set; available to download at the PredictDB Data Repository) from 10 brain tissues: anterior cingulate cortex, caudate basal ganglia, cerebellar hemisphere, cerebellum, cortex, frontal cortex, hippocampus, hypothalamus, nucleus accumbens basal ganglia, and putamen basal ganglia. We used a transcriptome-wide significant threshold of *p* < 1.34 × 10^-6^, which is the Bonferroni corrected threshold when adjusting for all tissues and genes (37,281 gene-based tests).

*Genetic correlations*: Genetic correlations between loneliness and 60 other traits were computed in LDSC.^31^ Here, the genetic correlation between traits is based on the estimated slope from the regression of the product of z-scores from two GWASs on the LD score and represents the genetic covariation between the two traits based on all polygenic effects captured by the included SNPs. Summary statistics from well-powered GWASs were available for 60 traits related to personality, cognition, reproduction, social circle, body composition, substance use, and physical and mental health. Multiple testing was corrected for using a Bonferroni corrected significance threshold of 8.5 × 10^-4^. LD scores were based on European populations from the HapMap 3 reference panel.^23,31^

*Polygenic scores:* All SNPs from the loneliness meta-analysis were thinned using an association-driven pruning algorithm that clumped SNPs into 250 kb windows and removed SNPs in LD (r2>0.1) with the most associated SNP (i.e., lowest p-value) in that window. LD estimates were directly derived from the BioVU samples (see below). After clumping, a total of 93,501 LD-independent SNPs remained for scoring. Scores were then constructed using PRSice software^32^ and defined by the sum of the number of risk alleles at each locus, weighted by their estimated effect sizes. The polygenic scores were calculated in an independent sample of 18,498 genotyped individuals of European descent in BioVU. Genotyping and QC of this sample have been described elsewhere.^20,33^

*PheWAS*: A logistic regression model was fitted to each of 897 case/control phenotypes to estimate the odds of each diagnosis given the loneliness polygenic score, after adjustment for sex, median age across the EHR, top 10 principal components of ancestry, and genotyping batch. The 897 disease phenotypes included 32 infectious diseases, 75 neoplasms, 86 endocrine/metabolic diseases, 29 hematopoietic diseases, 36 mental disorders, 44 neurological disorders, 54 sense organs, 126 circulatory system disorders, 59 respiratory diseases, 85 digestive diseases, 77 genitourinary diseases, 3 pregnancy complications, 43 dermatologic disorders, 64 musculoskeletal disorders, 8 congenital anomalies, 24 symptoms, and 52 injuries/poisonings. We required the presence of at least two International Classification of Disease (ICD) codes that mapped to a PheWAS disease category (Phecode Map 1.2 (https://phewascatalog.org/phecodes) to assign “case” status. PheWAS analyses were run using the PheWAS R package.^34^

*Lipid traits in the EHR*: We examined the relationship between polygenic risk for loneliness and three quantitative lipid traits. Clinically measured lipid levels included low density lipoprotein (LDL) (N = 6,455 with pre-medication values), high density lipoprotein (HDL) (N = 10,722), and triglycerides (trigs) (N = 11,012; Supplementary Table 5). As most patients had multiple lipid values available in their EHRs, we calculated median LDL, HDL, and triglyceride values for each patient after removing outlier values that were +/- 4 SDs from the sample mean. To adjust for age, we extracted the age at the median lipid value, and used the average age between the two median measurements if the number of lab value measurements was even. We then regressed the median lab value on sex and the cubic spline of median age, and quantile normalized the residuals. For sensitivity analyses, we also calculated the median of pre-medication (Supplementary Table 5) lipid values, using only observations that occurred before the first mention of lipid-lowering medication in the EHR,^35^ and transformed the age- and sex-adjusted residuals as above. Linear regression models were then fitted to the median LDL, HDL, and trigs values respectively to estimate the effect of the loneliness polygenic score on each lipid trait. As the lipid traits were already sex and age adjusted, we included only the top 10 principal components of ancestry and genotyping batch as covariates.

*Mendelian Randomization*: We performed two-sample bi-directional Mendelian Randomization (MR)^36^ analyses to investigate the direction of causality in the relationship between loneliness and cardiovascular risk factors and diseases. Of the eight cardiovascular risk factors and diseases for which we know the genetic correlations from the LDSC analyses (coronary artery disease [CAD], myocardial Infarction, high density lipoprotein (HDL) cholesterol, low density lipoprotein (LDL) cholesterol, total cholesterol, triglycerides, BMI, and body fat), we tested the four traits that showed a significant genetic correlation, namely CAD (*r_g_* = .19), triglycerides (*r_g_* = .14), BMI (*r_g_* = .18), and body fat (*r_g_* = .25). We used genome-wide significant SNPs from the five GWASs (loneliness and the four significant traits) to serve as instrumental variables (gene-exposure association). SNPs were pruned for LD (r^2^<.001), and the remaining SNPs (or proxy SNPs with r^2^≥0.8 when the top-SNP was not available in the other GWASs) were then identified in the GWAS summary statistics of the outcome variable (gene-outcome association). When both gene-exposure and gene-outcome associations are significant and in the expected ratio of an indirect causal effect, and the MR assumptions are met,^37^ this is considered evidence for a causal relationship. We combined estimates from individual SNPs by applying inverse-variance weighted (IVW) linear regression.^38^ We conducted three sensitivity analyses more robust to horizontal pleiotropy, each relying on distinct assumptions: weighted median regression^39^, MR-Egger regression^40^ and Generalized Summary-data based Mendelian Randomization (GSMR).^41^ Weighted median regression can provide a consistent estimate of a possible causal effect, even when up to 50% of the weight in the genetic instrument comes from invalid instruments. MR-Egger regression uses “Egger’s test” to test for bias from horizontal pleiotropy. MR-Egger will provide a consistent estimate of the causal effect, given that the strength of the genetic instrument (gene-exposure association) does not correlate with the effect that the instrument has on the outcome. This InSIDE assumption (Instrument Strength Independent of Direct Effect) is a much weaker assumption than the assumption that there is no pleiotropy. However, if the NOME (NO Measurement Error) assumption is violated, MR-Egger may be biased. Violation of NOME can be assessed with the I^2^ statistic, which ranges between 0 and 1. When I^2^ is below 0.9, there is a considerable risk of bias. By applying MR-Egger simulation extrapolation (SIMEX),^42^ this bias can be corrected for. When I^2^ is below 0.6 the results of MR-Egger (even with SIMEX correction) are not reliable. For our analyses we report MR-Egger results when I^2^ >0.9, MR-Egger SIMEX results when I^2^ = 0.6-0.9 and we don’t report MR-Egger results when I^2^ <0.6. Lastly, we performed GSMR, a method that takes into account LD between the different genetic variants included in an instrument. Since GSMR accounts for LD, we pruned the genetic variants included in GSMR instruments at a higher threshold of r^2^<0.05 (as opposed to r^2^<0.001). Including SNPs in higher LD than 0.05 was shown to provide very limited increase in power. GSMR includes a filtering step which excludes SNPs that are suspected to have pleiotropic effects on both the exposure and the outcome (HEIDI filtering).

## RESULTS

### GWAS Meta-Analysis

The proportion of phenotypic variance accounted for by all genotyped variants (SNP heritability) of the categorical loneliness measure in UKB and continuous loneliness measure were 13.3% (SE = .7) and 4.9% (SE = .8) respectively (see Supplementary Table 3). The results from the two meta-analyses (categorical and continuous phenotypes) were then meta-analyzed together using sample size-based weights.^24^ The adjusted effective sample size of the final meta-analysis, accounting for information from related individuals, was 205,708 (see Demontis et al).^24^ The SNP heritability of the final meta-analysis was 7.9% (SE = .4), which accounts for approximately one quarter of the total heritability as estimated in twin-family studies and is about twice as large as the SNP heritability estimate of a recently published GWAS on loneliness in the UK Biobank sample.^15^

The genomic inflation factor λ was 1.28 for the full meta-analysis (Figure 1) and results from LDSC analysis^23^ showed that this inflation was mostly due to true polygenic signal with about 1.8% of the inflation due to residual population stratification (LDSC intercept = 1.005, SE = .01, ratio = .015). We identified 14 independent genome-wide significant variants (r^2^ < .1), which were located in 12 genomic regions (i.e., within 250 kb; Figure 1, Table 1, and Supplementary File 1).

**Figure 1:**
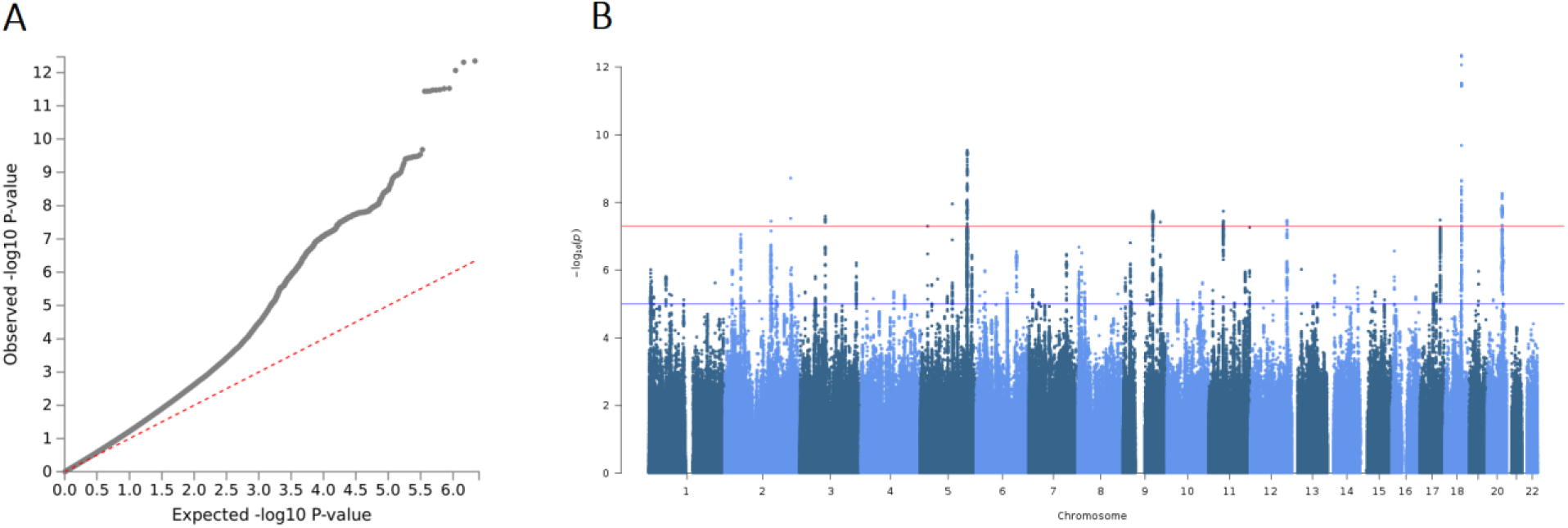
QQ-plot and Manhattan plot of meta-analysis on loneliness. **A**: The QQ-plot shows a considerable inflation of association statistics (λ = 1.28), which is mostly due to true polygenic signal rather than population stratification (LD-score regression intercept = 1.005, SE = .01, ratio = .015). **B:**Manhattan Plot of the Loneliness GWAS meta-analysis showing 14 independent genome-wide significant associations

**Table 1:** 14 Independent genome-wide significant SNPs from 12 loci, with independence based on an r2 threshold of .1, belonging to the same locus if they are within 250 kb (see supplementary file 1 for more details on the significant SNPs).

| SNPs | CHR | BP (hg19) | A1/A2 | BETA (SE) | p-value | Gene | Additional traits associated with top SNPs |
| --- | --- | --- | --- | --- | --- | --- | --- |
| rs618869;<br>rs12458015;<br>rs72627233 | 18 | 53248151;<br>53305735;<br>53486724 | C/T;<br>C/T;<br>G/T | .03 (.004);<br>.02 (.003);<br>.02 (.004) | $4.47 \times 10^{-13}$;<br>$2.35 \times 10^{-9}$;<br>$2.81 \times 10^{-8}$ | TCF4;<br>RPL21P126 | Schizophrenia, Autism, Fuchs's corneal dystrophy, Hand grip strength |
| rs4509081 | 5 | 152257172 | G/A | .02 (.003) | $2.90 \times 10^{-10}$ | LINC01470 | Life satisfaction, Bipolar disorder, Schizophrenia |
| rs74338595 | 2 | 212749786 | C/T | -.02 (.003) | $1.92 \times 10^{-9}$ | ERBB4 | |
| rs1022688 | 20 | 47648856 | A/G | .02 (.003) | $5.41 \times 10^{-9}$ | ARFGEF2 | Subjective well-being, Positive affect, Height, Cognitive ability |
| rs171697 | 5 | 103956516 | G/C | .02 (.003) | $1.10 \times 10^{-8}$ | AC099520.1 | Anorexia nervosa, Depression |
| rs10123378 | 9 | 96375217 | C/T | -.02 (.003) | $1.79 \times 10^{-8}$ | PHF2 | Childhood BMI, Educational attainment |
| rs7925389 | 11 | 47466790 | T/A | -.02 (.003) | $1.81 \times 10^{-8}$ | RAPSN | Subjective well-being, Neuroticism, Height, BMI |
| rs7626596 | 3 | 82000680 | A/G | -.02 (.003) | $2.55 \times 10^{-8}$ | | |
| rs11867618 | 17 | 65875587 | A/G | .02 (.004) | $3.29 \times 10^{-8}$ | BPTF | BMI, Eosinophil and basophil counts, Lung cancer, Alcohol dependence |
| rs61943369 | 12 | 118805950 | T/C | .02 (.004) | $3.35 \times 10^{-8}$ | TAOK3 | Glucose homeostasis traits, Neuroticism |
| rs2656321 | 2 | 149073434 | T/C | .02 (.003) | $3.57 \times 10^{-8}$ | MBD5 | |
| rs2149351 | 9 | 120501644 | G/T | -.02 (.004) | $3.78 \times 10^{-8}$ | | Neuroticism |

### MAGMA and S-PrediXcan gene-based analyses

We performed two types of gene-based analyses using MAGMA, which aggregates SNP effects at the gene level using positional annotations, and S-PrediXcan, which uses expression quantitative-trait loci (eQTL) annotations to assign SNPs to genes. The meta-analysis summary statistics formed the basis to compute gene-based *p*-values in MAGMA^25^ and S-PrediXcan^29^ for 17,715 and 13,037 protein coding genes, respectively. In the MAGMA analysis, a total of 38 genes reached genome-wide significance at a Bonferroni corrected significance threshold of 2.82 × 10^-6^ (Supplementary Figure 1). Six genes of these genes (*ARFGEF2*, *BPTF*, *MBD5*, *PHF2*, *TAOK3*, *TCF4*; Table 1) included a genome-wide significant SNP from the GWAS meta-analysis. Using S-PrediXcan,^29^ we identified 10 genes (of which 8 were significant in the MAGMA analysis) that significantly associated with loneliness at a Bonferroni corrected significance threshold of *p* < 1.34 x 10^-6^ across six brain tissues: anterior cingulate cortex, cerebellar hemisphere, cerebellum, prefrontal cortex, cortex, and the caudate basal ganglia (Supplementary Table 4).

### GWAS signals are significantly enriched for brain tissues and evolutionary conserved regions

Next, we investigated if genetic effects on loneliness were enriched for loci with specific functional and tissue annotations.

First, we tested whether genome-wide effects on loneliness were consistent with tissue-specific differential gene-expression based on GTEx RNA-sequence data from 53 tissues types using two approaches. For the first approach, we determined whether the distribution of effect sizes of all 17,715 protein coding genes estimated from the gene-based tests showed enrichment of expression across multiple tissues.^26^ These results indicated that the gene-based association results were significantly enriched (Bonferroni threshold: *p* < 9.4 × 10^-4^) for genes with higher gene-expression levels in five brain tissues: frontal cortex, cortex, cerebellar hemisphere, cerebellum, and anterior cingulate cortex (Figure 3). For the second approach, SNP-heritability of loneliness was partitioned into categories of functional SNP annotations using LDSC.^23^ We found that SNPs associated with loneliness were also more likely than expected by chance (Bonferroni threshold: *p* < 9.4 × 10^-4^) to regulate gene expression in four brain tissues including cerebellum, anterior cingulate cortex, substantia nigra, and cortex.

**Figure 3:**
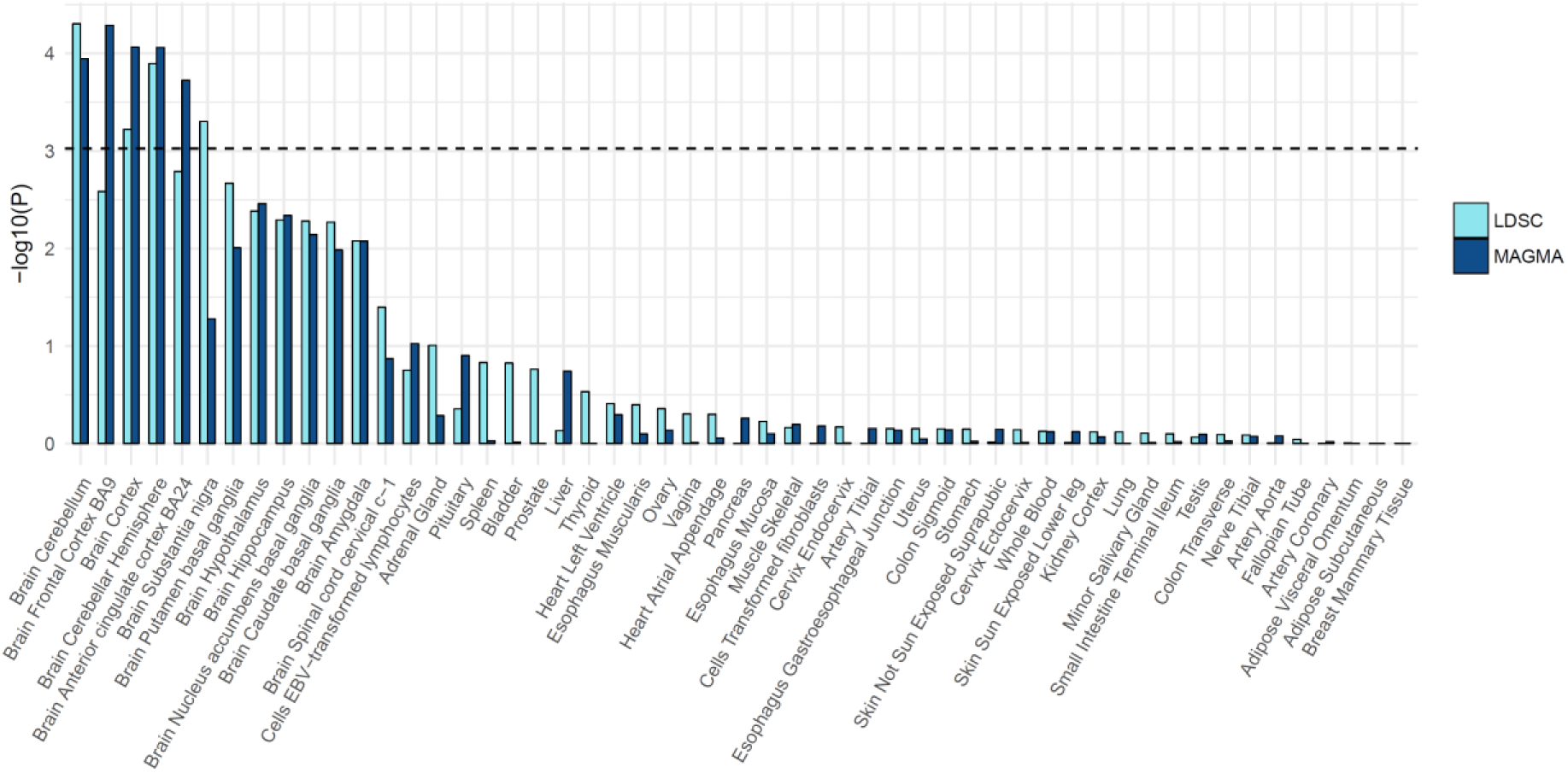
Enrichment of gene-expression for 53 specific tissue types using MAGMA and LDscore regression.

Second, we used LDSC to test for the enrichment of 24 genomic annotations that are not specific to any cell type, including coding vs non-coding regions, promoter regions, introns, and evolutionary conserved regions (see Finucane et al, 2015^28^ for additional details). Of these 24 annotations, the genetic signals were significantly enriched for regions that were highly evolutionary conserved in mammals, which contain 2.6% of all SNPs but explain 25% of the loneliness heritability captured by all SNPs (Figure 4).

**Figure 4:**
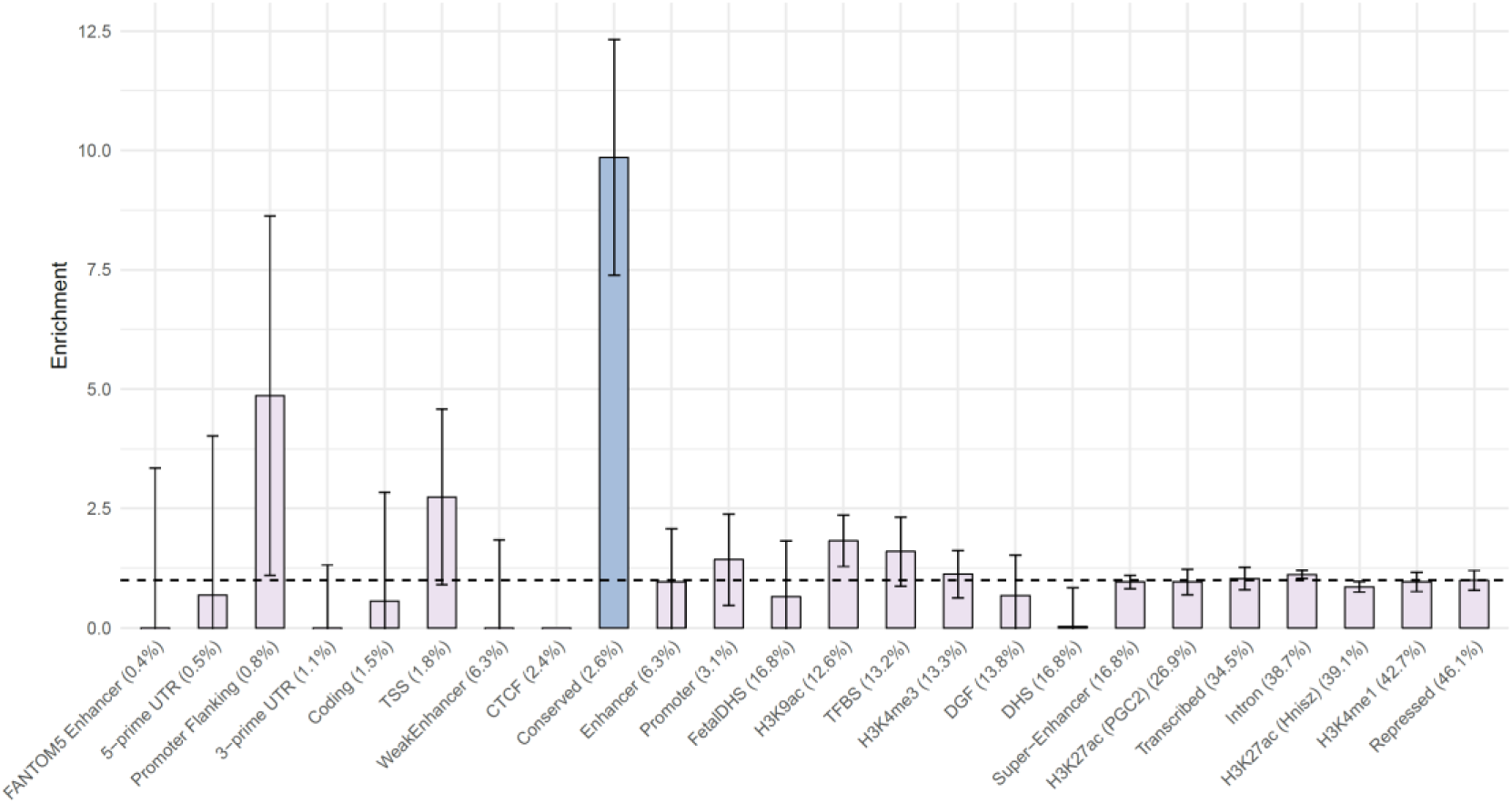
Enrichment of 24 annotations not specific to cell-types, ordered by size (proportion of SNPs).

### Genetic Correlations

Genetic correlations^31^ were estimated for loneliness and 60 characteristics from 9 domains including anthropomorphic traits, cardiovascular disease risk, cognitive functions, mental health, reproduction, and substance use. After applying a Bonferroni corrected significance threshold of 8.3 × 10^-4^, 34 out of 60 traits showed a significant genetic correlation with loneliness (Figure 5 & supplementary_file.csv). A significant signal was observed at least once from each of the 9 domains, with the strongest genetic correlations observed for mental health, especially for depressive symptoms (*r_g_* = .88, *p* = 2.2 × 10^-101^), subjective wellbeing (*r_g_* = −.77, *p* = 8.0 × 10^-50^), and major depressive disorder (*r*_g_ = .64, *p* = 2.8 × 10^-114^). In the health domain, tiredness and self-rated health showed the strongest correlations (*r_g_* = .74, *p* = 3.0 × 10^-59^, and *r_g_* = −.56, *p* = 2.5 × 10^-44^, respectively; more loneliness = more tiredness and worse health), while father’s and mother’s age of death showed modest but significant negative genetic correlations with loneliness (*r_g_* = −.32, *p* = 1.8 × 10^-5^, and *r_g_* = −.38, *p* = 1.8 × 10^-5^, respectively). Four out of five personality dimensions showed a significant genetic correlation with loneliness, with neuroticism showing the highest association (*r_g_* = .69, *p* = 2.4 × 10^-49^), a genetic association that has recently been shown to be a major driver for the association between loneliness and personality.^14^ SES indicators related to economic success (Townsend index and income; *r_g_* = .43, *p* = 7.7 × 10^-12^, and *r_g_* = −.50, *p* = 3.5 × 10^-51^, respectively) showed a considerably higher genetic correlation with loneliness than indicators of cognition (IQ and educational attainment; *r_g_* = −.19, *p* = 6.1 × 10^-6^, and *r_g_* = −.28, *p* = 3.7 × 10^-23^, respectively). Genetic correlations with traits from the reproduction domain indicate that having more offspring and having offspring at a younger age is genetically associated with higher levels of loneliness, an association that is in the other direction for phenotypic correlations.^12^ For substance use, alcohol consumption had a significant genetic correlation with loneliness (*r_g_* = −.16, *p* = 4.9 × 10^-4^), with more alcohol consumption being associated with lower loneliness, while alcohol dependence had a larger genetic correlation in the opposite direction (*r_g_* =.43, *p* = 9.7 × 10^-7^). In the social circle domain, family and friendship satisfaction both showed significantly larger genetic correlations with loneliness (*r_g_* = .56, *p* = 2.9 × 10^-42^, and *r_g_* = .55, *p* = 4.6 × 10^-31^, respectively; more loneliness = less satisfaction) than the frequency of friend and family visits (*r_g_* = .18, *p* = 9.8 × 10^-6^; more loneliness = less visits), suggesting that the subjective experience of social isolation may play a larger role in feeling lonely than objective social isolation.

**Figure 5:**
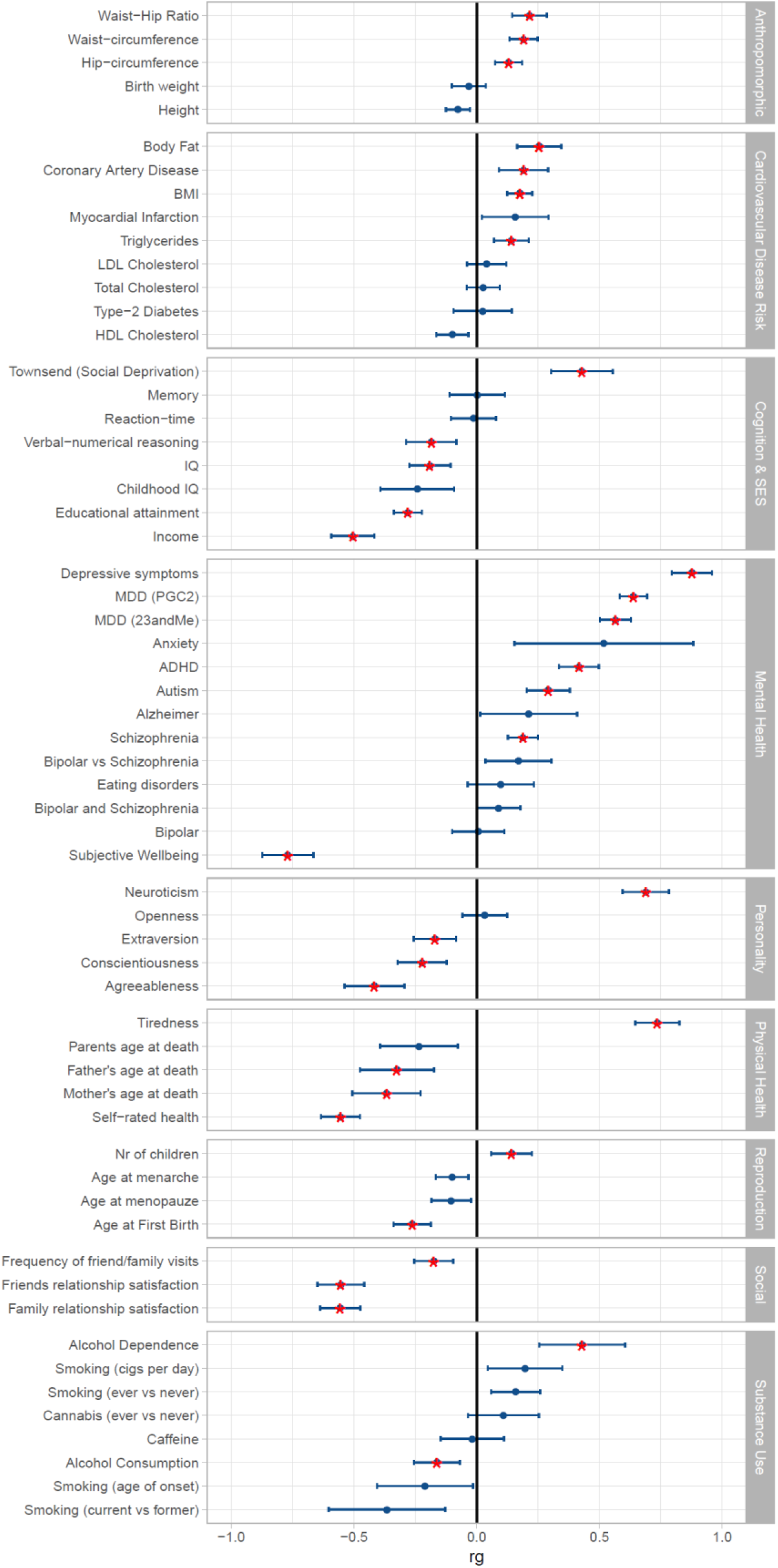
Genetic correlations as computed with LD score regression. Red stars are significant after Bonferroni correction.

### PheWAS on the polygenic score for loneliness

Five cardiovascular, three neuropsychiatric, and one of the metabolic phenotypes were significantly associated with genetic propensity to loneliness after Bonferroni correction for the 897 phenotypes tested (*p* < 5.57 × 10^-5^) (Figure 6). Mood disorders yielded the most significant association with the loneliness polygenic score (N_cases_ = 3,299, OR = 1.11, SE = .02, *p* = 2.8× 10^-7^), followed by depression (N_cases_ = 2,969, OR = 1.11, SE =.02 *p* = 3.9 × 10^-7^), heart failure (N_cases_ = 818, OR = 1.19, SE = .03, p = 4.9 x 10^-6^), ischemic heart disease (N_cases_ = 5,797 OR = 1.09, SE = .02, *p* = 5.7 × 10^-7^), and tobacco use disorder (N_cases_ = 1,705, OR = 1.12, SE = .03, *p* = 1.2 × 10^-5^). Complete results may be viewed interactively at https://sealockj.shinyapps.io/loneliness_phewas/loneliness_phewas.Rmd.

**Figure 6:**
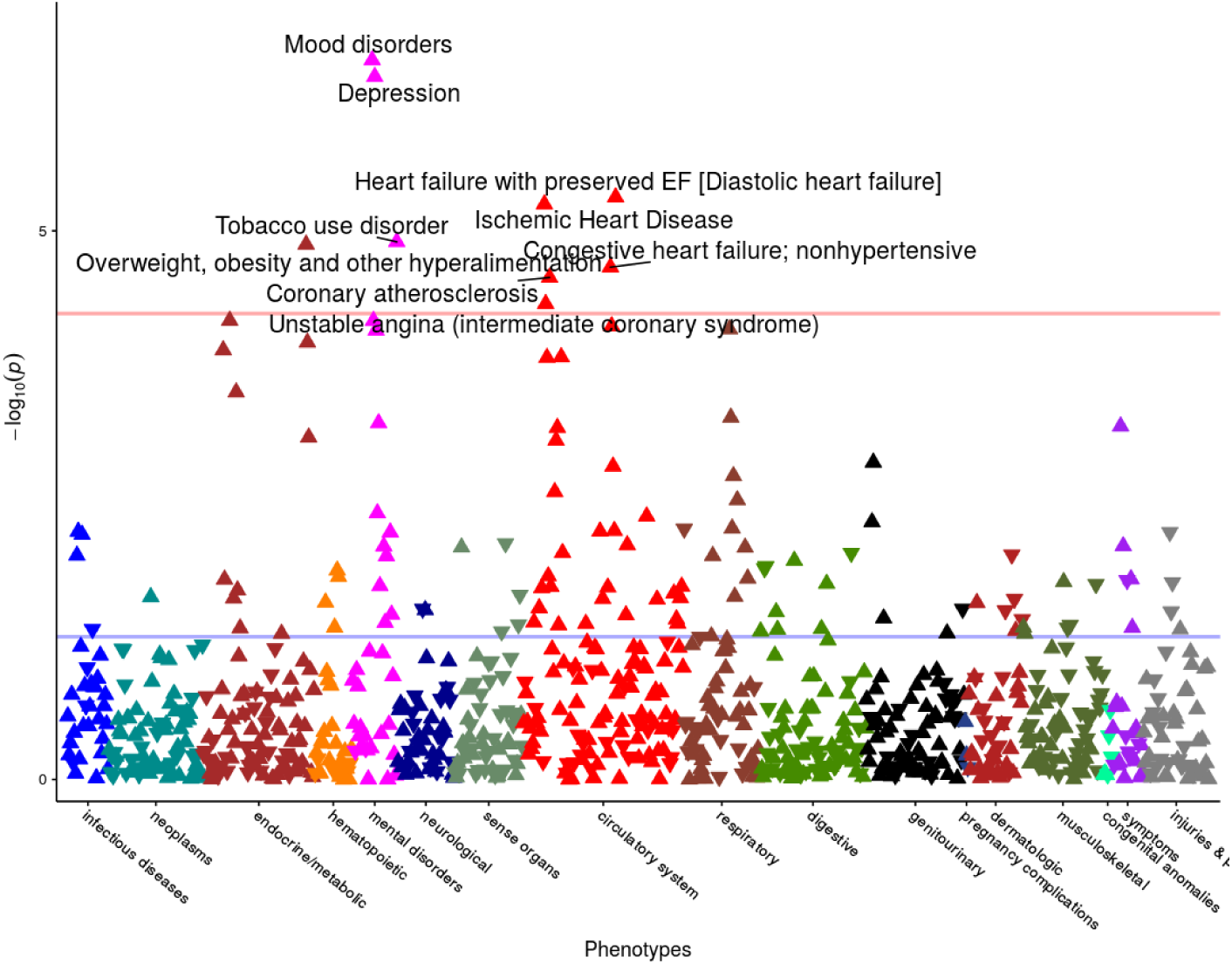
Results of the Phewas on the polygenic score for loneliness, corrected for gender, age, first 10 PCs, and batch.

In our subsequent analysis of quantitative lipid traits, the polygenic score for loneliness modestly but significantly predicted reduced HDL (R^2^ = 0.16%, *p* = 2.99 x 10^-5^) and increased triglycerides (R^2^ = 0.16%, *p* = 2.40 x 10^-5^), but not LDL levels (R^2^ = 0.05%, *p* = 2.62 x 10^-2^). To benchmark these results, we compared them to the proportion of variance explained by a polygenic score for CAD developed using the beta weights from the CARDIOoGRAMplusC4D study (http://www.cardiogramplusc4d.org/data-downloads/).^43^ The proportion of variance explained by the polygenic score for CAD was similar in magnitude to the variance explained by the loneliness polygenic score for clinically evaluated HDL (R^2^ = 0.34%, *p* = 6.96 x 10^-10^), triglycerides (R^2^ = 0.16%, *p* = 2.40 x 10^-5^), and LDL (R^2^ = 0.03%, *p* = 6.05 x 10^-2^). As a negative control (see Supplementary Figure 3), we also tested whether the loneliness polygenic score predicted median height across the medical record and, as expected, observed no significant prediction of height (R^2^ = 0.02%, *p* = .09) (Supplementary Figure 3). Details on the best-fit p-value thresholds used in these analyses are provided in Supplementary Table 7 and Supplementary Figure 4.

### Mendelian Randomization

To distinguish genetic correlation from causation, we applied Mendelian Randomization (MR; methodology to examine evidence for causal effects of one phenotype on another) to traits that showed a significant genetic correlation with loneliness and of which the top SNPs were unlikely to share pleiotropic effects with loneliness. We focused on the relationship between loneliness and cardiovascular disease and its associated risk factors including: coronary artery disease (CAD), myocardial infarction, HDL cholesterol, LDL cholesterol, total cholesterol, triglycerides, BMI, and body fat, which we also found to be genetically correlated with loneliness. Of these traits, we selected those with a significant genetic correlation with loneliness, namely CAD (*r_g_* = .19), triglycerides (*r_g_* = .14), BMI (*r_g_* = .18), and body fat (*r_g_* = .25) (see Figure 7). When both gene-exposure and gene-outcome associations are significant and in the expected ratio of a causal effect, and the MR assumptions are met,^37^ this is considered evidence for a causal relationship.

**Figure 7:**
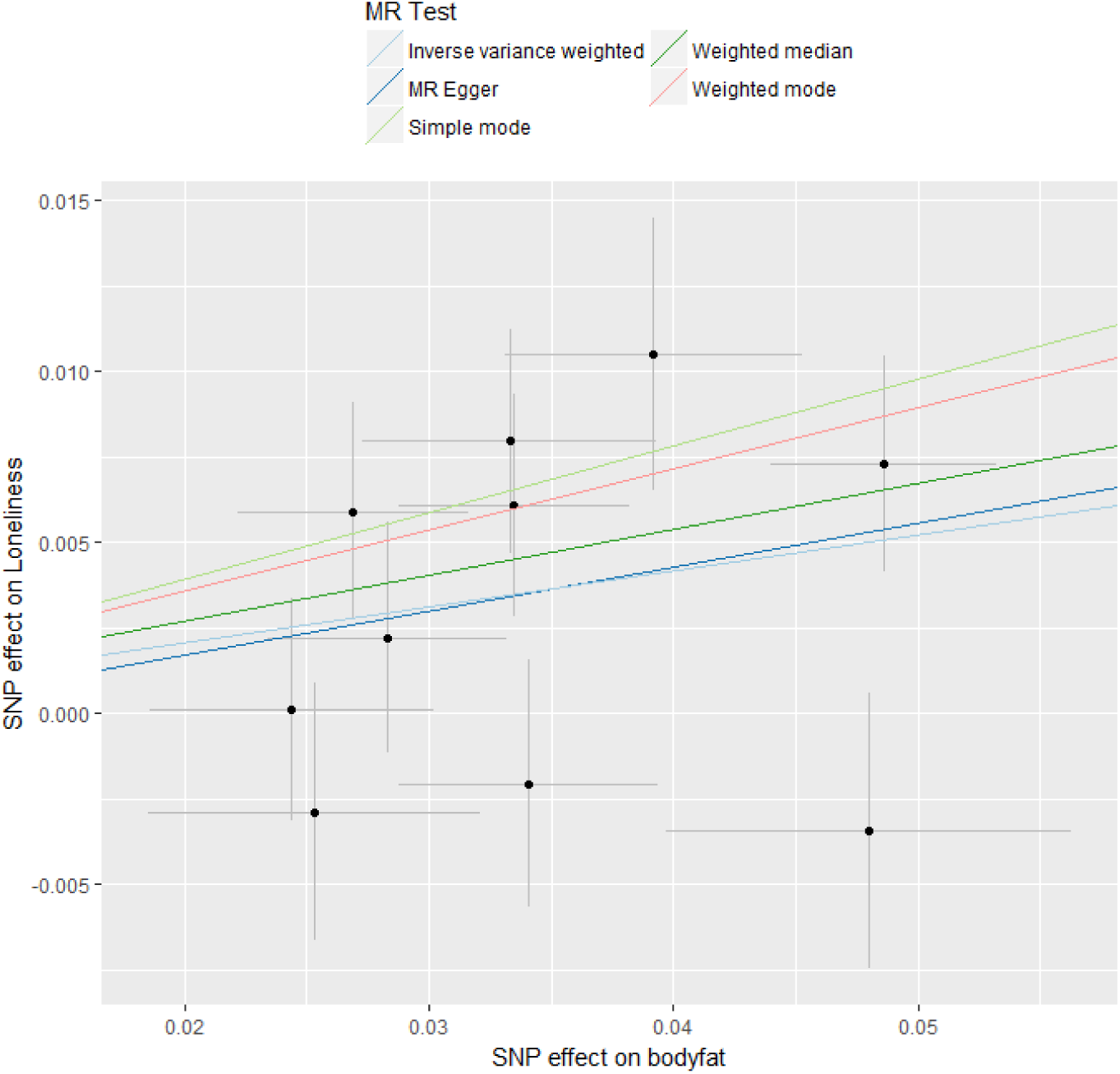
Two Sample Mendelian Randomization results for the causal effect of body fat on loneliness.

We found evidence for a causal effect of body fat on loneliness using the IVW, weighted median and GSMR methods, but not the MR-Egger method (higher body fat = higher loneliness; Table 2 and Figure 7). Since the effect size of MR-Egger is of similar magnitude to the other two analyses and the MR Egger intercept is not significantly different from 0 (*p* = .90, see Supplementary Table 12), indicating that there is no evidence of horizontal pleiotropy for this relationship, it is most likely that this analysis remained underpowered due to weak instrument variables.^44^ There was also evidence for a causal, increasing effect of BMI on loneliness, but only with the GSMR method. We note that sample overlap between GWASs could cause a bias of MR results in the direction of the observational association. However, sample overlap was minimal in the present study (max 3.7%) and so it is unlikely to have affected our findings.

**Table 2.** Two sample, bidirectional Mendelian randomization results

| Exposure | Outcome | *n* | IVW |  |  | Weighted<br>median |  |  | MR-Egger |  |  | GSMR |  |  |  |
| --- | --- | --- | --- | --- | --- | --- | --- | --- | --- | --- | --- | --- | --- | --- | --- |
|  |  | SNPs | beta | *SE* | *P* | Beta | *SE* | *p* | beta | *SE* | *p* | SNPs | Beta | *SE* | *p* |
| Loneliness | BMI | 9 | -.23 | .22 | .30 | .04 | .13 | .73 | .88* | 1.72 | .62 | 8 | .02 | .13 | .87 |
| Loneliness | Body fat | 9 | .00 | .20 | .997 | .05 | .13 | .73 | .58* | 1.71 | .74 | 8 | .25 | .17 | .14 |
| Loneliness | Triglycerides | 9 | -.06 | .17 | .72 | .02 | .14 | .90 | *n.a.* | *n.a.* | *n.a.* | 9 | -.09 | .14 | .53 |
| Loneliness | CAD | 12 | -.01 | .19 | .97 | .24 | .22 | .27 | *n.a.* | *n.a.* | *n.a.* | 13 | .14 | .23 | .54 |
| BMI | Loneliness | 53 | .03 | .03 | .22 | .02 | .03 | .59 | .04 | .07 | .61 | 65 | **.06** | **.02** | **.004** |
| Body fat | Loneliness | 10 | **.10** | **.04** | **.01** | **.14** | **.05** | **.003** | .17* | .25 | .52 | 10 | **.12** | **.04** | **.001** |
| Triglycerides | Loneliness | 41 | .00 | .01 | .85 | -.01 | .02 | .75 | -.01 | .02 | .77 | 77 | .00 | .01 | 0.86 |
| CAD | Loneliness | 26 | .00 | .01 | .76 | .01 | .01 | .50 | -.02* | .02 | 0.29 | 31 | .03 | .02 | .20 |

\*Egger-SIMEX correction applied because of low $I^2$ estimates. n.a. = $I^2$ estimates too low to give reliable results for MR-Egger.

## Discussion

Chronic loneliness is strongly associated with physical and mental health, and is a growing concern in many societies. In this study we performed a large GWAS meta-analysis which confirmed the polygenic architecture of loneliness, and performed follow-up analyses to further investigate the genetic architecture of loneliness and its relationship with a wide range of traits. We identified 14 SNPs located in 12 independent loci that were significantly associated with loneliness. The most significant SNP signal came from chromosome 18 within the *TCF4* gene, which plays an important role in nervous system development and has recently been associated with major depressive disorder (MDD)^45^. We replicated 10 loci from a previous GWAS on loneliness^15^ and report two novel loci on chromosomes 9 and 12 (top SNPs: rs2149351 and rs61943369, respectively).

We identified that the significantly associated variants were disproportionally enriched for regions that were conserved in mammals, confirming the biological importance of processes that underlie individual differences in loneliness. Additionally, we found that associated variants were highly enriched in the brain, in particular in the cerebellum, (frontal) cortex, anterior cingulate cortex, and substantia nigra. The cerebellum is mostly known for its modulating role in motor, cognitive, and affective functions, and has been shown to play a role in social cognition as well, especially for processes that require higher-level abstraction away from the current event (i.e., past, future or hypothetical events).^46^ The prefrontal cortex is posited to be involved in the perception of social isolation (i.e., loneliness).^47-49^ The anterior cingulate cortex is functionally connected with the prefrontal cortex in orchestrating emotional and physiological adjustments for potential threats and stressors, and is known to be involved the social (rather than the physical) pain associated with loneliness.^50^ The substantia nigra is most known for its role in reward and learning, which extends to social contexts as well.^51^ A recent large GWAS meta-analysis on MDD performed similar tissue enrichment analyses and reached only partly the same conclusions: for MDD all the same cortical regions were significant (frontal cortex, cortex, anterior cingulate cortex, substantia nigra), but none of the cerebellar regions (cerebellar hemisphere and cerebellum).^45^ This suggests that the role of the cerebellar region may be more specific or larger for loneliness. The complex processes underlying loneliness likely involve more brain regions, as all 13 brain regions included in the enrichment analyses depicted in Figure 3 were nominally significant with *p* < .05.

Over 34 out of the 60 traits in the genetic correlation analyses showed a significant genetic association with loneliness, suggesting widespread shared genetic influences (e.g., pleiotropic effects) or causal relationships between loneliness and several traits. MDD has been strongly associated with loneliness in previous studies^52-54^, but evidence from longitudinal studies indicates that loneliness and depression are conceptually and statistically different constructs.^53-55^ Our results confirm the strong biological ties between loneliness, major depression, and depressive symptoms in both research ascertained samples and EHR from a hospital population. Our analyses do not provide conclusive findings however regarding the direction of causation in the relationship between loneliness and MDD, due to a lack of instrument variables for loneliness that are strong enough for causal inference.

There are several traits that show a genetic correlation with loneliness that is not in line with the expected direction from phenotypic observational data. In observational data, having more offspring has been shown to be associated with lower levels of loneliness,^12^ while the genetic correlations with number of offspring and age at first birth indicate an association with loneliness in the opposite direction (i.e., having more children or children at an earlier age = more lonely). Alcohol use has also been associated with higher levels of loneliness^56^, while we find a genetic correlation with alcohol consumption in the opposite direction. A possible explanation of these contradictory results may be that they are driven by the genetic association between loneliness and socio-economic status, which is also associated with many life outcomes. We observe a significant genetic overlap between loneliness and SES-related traits (e.g., income, educational attainment [EA], social deprivation of the neighborhood), with lower SES indicators showing a genetic association with more loneliness (with a particularly strong genetic correlation for income: *r*_g_ = −.50). Number of offspring shows a negative genetic correlation with EA^57^ (and positive with loneliness), age at first birth a positive genetic correlation with EA^57^ (and negative with loneliness), while alcohol consumption shows a positive genetic correlation with EA^58^ (and negative with loneliness), and alcohol dependence a negative genetic correlation with EA^59^ (and positive with loneliness), which is all in line with the negative genetic association between loneliness and EA/SES.

Our phenome-wide analysis in a unique EHR dataset recapitulated the genetic correlation results and found that genetic propensity to loneliness is associated with increased risk for clinical depression, cardiovascular disease, and metabolic diseases such as type-2 diabetes. Furthermore, we found that elevated triglycerides and reduced HDL, two well-known risk factors for heart disease, were also associated with predisposition to loneliness after adjusting for covariates and even after restricting to levels prior to use of antilipemic medications. These findings provide a proof of principle that even in clinical settings, polygenic scores can be used to uncover relationships between difficult to measure behavioral traits (such as loneliness) and health outcomes. Another important advantage of this out-of-sample analysis is that by relying on physician assigned ICD codes instead of retrospective self-report (as in UK Biobank), we avoid potential reporting biases related to loneliness that may subsequently influence correlations between loneliness and health outcomes. One possible limitation of this out-of-sample analysis could be the potential for some overfitting because we included all SNPs at a p-value threshold of 1. However, this is unlikely to be a substantial driver of our results given that an analysis of loneliness polygenic scores using a p-value threshold of 0.05 yielded similar results (Supplementary Figure 5). Another important limitation is that polygenic score analyses cannot distinguish pleiotropy from causal effects.

With Mendelian randomization analysis – which can distinguish pleiotropic from causal effects – we found evidence of a causal, increasing, effect of body fat on loneliness, and weaker evidence of a causal, increasing, effect of BMI on loneliness. This concurs with a recent MR study reporting that BMI increases depressive symptoms and decreases subjective well-being.^60^ Our evidence for a causal effect on loneliness was stronger for total body fat than for BMI, which may be due to body fat being a better measure for an unhealthy excess of body weight than BMI. Nonetheless, both findings point to an increased body weight causally leading to poorer mental health. Our MR analysis ruled out the possibility of horizontal pleiotropy among the instrument variables used in this analysis. However, it is important to note that the condition of “no pleiotropy” is only required for the instrument variables themselves and need not apply genome-wide. Indeed, it is possible (and likely) that the relationship between loneliness and health outcomes is influenced by bidirectional causal effects and pleiotropic biological effects. More research is needed to tease apart the complex etiology of these states and traits.

We identified 12 genome-wide significant loci and a total of 40 significantly associated genes. Follow-up analyses identified specific brain tissues in cortical and cerebellar regions involved in loneliness risk. We showed that a wide range of traits are genetically associated with loneliness and extended these findings to an electronic health record system. Limitations of the study include the sample size, which, while large even by modern GWAS standards is still modest for detection of genome-wide significant loci for loneliness given its heritability and heterogeneity. Future work needs to establish the etiology of these associations, and to determine which additional loci explain the rest of common genetic variation underlying loneliness, which together explain ~8% of individual differences.

## ACKNOWLEDGEMENTS AND FUNDING

### Acknowledgements

**We dedicate this paper to the memory of Dr. John T. Cacioppo, who pioneered the scientific study of loneliness**

We warmly thank all volunteer participants who contributed data to this project.

Preparation of this manuscript was supported by the National Institutes of Health (NIH, R01AG033590 to JC). NTR: DIB acknowledges the Royal Netherlands Academy of Science Professor Award (PAH/6635). Data collection and genotyping in NTR were supported by the Netherlands Organization for Scientific Research (904‐61‐090, 85‐10‐002,904‐61‐193,480‐04‐004, 400‐05‐717, Spi-56‐464‐14192 and 480-15-001/674); Biobanking and Biomolecular Resources Research Infrastructure (BBMRI –NL, 184.021.007 and 184.033.111); the Avera Institute for Human Genetics, Sioux Falls, South Dakota (USA) and the National Institutes of Health (NIH, R01D0042157‐01A), Grand Opportunity grants 1RC2MH089951‐01 and 1RC2 MH089995‐01 from the NIMH. JLT is supported by a Rubicon grant from the Netherlands Organization for Scientific Research (NWO; grant number 446-16-009). LKD is supported by a grant from the NIMH (5R01MH113362-02) and JMS is supported by an NIH training grant (2T32GM080178).

The dataset(s) used for the PheWAS analyses described were obtained from Vanderbilt University Medical Center’s BioVU which is supported by institutional funding, the 1S10RR025141-01 instrumentation award, and by the CTSA grant UL1TR000445 from NCATS/NIH. Additional funding provided by the NIH through grants P50GM115305 and U19HL065962. The authors wish to acknowledge the expert technical support of the VANTAGE and VANGARD core facilities, supported in part by the Vanderbilt-Ingram Cancer Center (P30 CA068485) and Vanderbilt Vision Center (P30 EY08126).

*Conflict of interests*: PF, SLE and members of the 23andMe research team are employees of 23andMe Inc.

*Data access:* The full summary statistics for the 23andMe dataset will be made available to qualified investigators who enter into an agreement with 23andMe that protects participant privacy. Interested investigators should visit research.23andMe/collaborate/#publication to learn more and to apply for access.

---

**Supplementary Table 1:** Subjects and phenotype details per cohort

| Dataset | Sample Size | % Females | Mean Age (SE) | Loneliness Measure |
| --- | --- | --- | --- | --- |
| UK Biobank<sup>61</sup> | 413,337 | 54% | 56.5 (8.1) | Do you often feel lonely? (yes/no) |
| 23andMe<sup>62</sup> | 20,591 | 55% | 59.6 (16.0) | 9-item questionnaire (4-point scale):<br>1) I feel in tune with the people around me<br>2) There are people I can turn to.<br>3) I feel alone<br>4) I feel part of a group of friends<br>5) I have a lot in common with the people around me<br>6) I feel isolated from others<br>7) There are people who really understand me<br>8) I am unhappy being so withdrawn<br>9) There are people I can talk to |
| Netherlands Twin Register<sup>63</sup> | 11,046 | 63% | 44.6 (17.1) | 3-item questionnaire (3-point scale):<br>1) How often do you feel left out?<br>2) How often do you feel isolated from others?<br>3) How often do you feel that you lack companionship? |
| Health & Retirement Study<sup>13</sup> | 7,556 | 59% | 67.2 (10.3) | 3-item questionnaire (3-point scale):<br>1) How often do you feel left out?<br>2) How often do you feel isolated from others?<br>3) How often do you feel that you lack companionship? |
| Rotterdam Study<sup>64</sup> | 7,764 | 57% | 67 (10.7) | Did you feel lonely during the past week? (4-point scale) |
| Sweden – SALTY<sup>65</sup> | 5,750 | 52.2% | 49.8 (4.2) | Did you feel lonely during the past week? (4-point scale) |
| Sweden – TwinGene<sup>65</sup> | 9,617 | 53% | 58.4 (8.0) | Did you feel lonely during the past week? (4-point scale) |

**Supplementary Table 2:** Genotyping, imputation, QC, and GWAS information per cohort

| Dataset | Microarray(s) | Imputation Panel | SNP MAF, INFO, HWE exclusion thresholds | N SNPs after QC | Identification of European Ancestry | GWAS analysis |
| --- | --- | --- | --- | --- | --- | --- |
| UK Biobank<sup>61</sup> | UK BiLEVE array & UK Biobank Axiom Array | Haplotype Reference Consortium | MAF <.005;<br>INFO <.8;<br>HWE $p < 10^{-10}$ | 7,954,332 | Self-report & PCA | Logistic regression in PLINK in 3 separate groups of unrelates: 332,991 + 57,865 British, and 22,481 of Non-British European (74,142 cases & 339,195 controls). Covariates: sex, age, 40 PCs. |
| 23andMe<sup>62</sup> | 23andMe custom genotyping array platforms<sup>62</sup> | 1000 Genomes (Phase 1 Version 3) | MAF <.01;<br>INFO <.5;<br>HWE $p < 10^{-20}$ | 14,113,458 | Ancestry Composition<sup>66</sup> | Linear regression using the internal 23andMe pipeline.<sup>62</sup> Covariates: age, sex, 5 PCs, genotype platform. |
| Netherlands Twin Register<sup>63</sup> | Several Illumina and Affymetrix platforms<sup>63,67</sup> | Haplotype Reference Consortium | MAF <.01;<br>INFO <.4;<br>HWE $p < 10^{-5}$ | 6,917,809 | PCA | PLINK linear regression with within family clustering. Covariates: sex, age, 10 PCs |
| Health & Retirement Study<sup>13</sup> | Illumina Human Omni-2.5 | Haplotype Reference Consortium | MAF <.01;<br>INFO <.5;<br>HWE $p < 10^{-6}$ | 5,768,559 | NA | Linear mixed model in GEMMA. Covariates: sex, age, marital status |
| Rotterdam Study<sup>64</sup> | Several Illumina and Affymetrix platforms<sup>64</sup> | Haplotype Reference Consortium | MAF <.05;<br>INFO <.4;<br>HWE $p < 10^{-7}$ | 6,984,254 | PCA | Linear mixed model in GCTA. Covariates: sex, age, 10 PCs |
| Sweden – SALTY<sup>65</sup> | Illumina Infinium PsychArray-24 | 1000 Genomes (Phase 1 Version 3) | MAF <.01;<br>INFO <.8;<br>HWE $p < 10^{-6}$ | 8,511,408 | PCA | Linear mixed model in RAREMETALWORKER. Covariates: sex, age, 10 PCs |
| Sweden – TwinGene<sup>65</sup> | Illumina OmniExpress | 1000 Genomes (Phase 1 Version 3) | MAF <.01;<br>INFO <.8;<br>HWE $p < 10^{-6}$ | 8,834,367 | PCA | PLINK linear regression with within family clustering. Covariates: sex, age, 10 PCs |

**Supplementary Table 3:** Sample size, lambda, intercept, and h2 estimated from LD score regression

|  | UKB<br>unrelateds | UKB<br>relatives | UKB NBW | 23andMe | HRS | NTR | Rotterdam | Sweden -<br>SALTY | Sweden -<br>TwinGene |
| --- | --- | --- | --- | --- | --- | --- | --- | --- | --- |
| *N* | 332,991 | 57,862 | 22,481 | 20,591 | 7,556 | 11,046 | 7,764 | 5,750 | 9,617 |
| *Lambda* | 1.233 | 1.053 | 1.011 | 1.047 | 1.017 | 1.002 | 1.002 | 1.011 | 1.011 |
| *Intercept* | 1.017<br>(.008) | 1.012<br>(.007) | 1.005<br>(.007) | 1.001<br>(.006) | 1.006<br>(.006) | 1.005<br>(.006) | .994 (.006) | .99 (.007) | 1.020<br>(.006) |
| *h<sup>2</sup>* | .081 (.01) | .091 (.02) | .026 (.04) | .096 (.02) | .106 (.06) | .002 (.04) | .031 (.06) | .108 (.08) | -.047 (.04) |

**Supplementary Table 4:** Significant associations of S-PrediXcan gene-based association analyses (Bonferroni corrected significance threshold = 0.05/37281 = 1.34 × 10^-6^)

| Gene | Gene name | Z score | Effect Size | p-value | Var g | Pred perf r2 | Pred perf pval | N SNPs used | N SNPs in cov | N snps in model | Tissue |
| --- | --- | --- | --- | --- | --- | --- | --- | --- | --- | --- | --- |
| ENSG00000249484 | AC091969.1 | 6.176 | 0.080 | 6.58E-10 | 0.030 | 0.112 | 1.25E-04 | 23 | 31 | 31 | caudate |
| ENSG00000196666 | FAM180B | -5.700 | -0.042 | 1.20E-08 | 0.074 | 0.076 | 1.53E-03 | 22 | 26 | 26 | cerebellum |
| ENSG00000165915 | SLC39A13 | -5.631 | -0.039 | 1.79E-08 | 0.092 | 0.168 | 1.77E-05 | 11 | 20 | 20 | prefrontal cortex |
| ENSG00000124198 | ARFGEF2 | -5.434 | -0.138 | 5.50E-08 | 0.006 | 0.105 | 5.31E-04 | 7 | 12 | 12 | cortex |
| ENSG00000109919 | MTCH2 | -5.420 | -0.050 | 5.96E-08 | 0.052 | 0.140 | 8.06E-05 | 23 | 30 | 30 | prefrontal cortex |
| ENSG00000196666 | FAM180B | -5.415 | -0.438 | 6.12E-08 | 0.001 | 0.140 | 1.24E-04 | 1 | 12 | 12 | prefrontal cortex |
| ENSG00000196666 | FAM180B | -5.415 | -1.961 | 6.12E-08 | 0.000 | 0.049 | 2.51E-02 | 1 | 13 | 13 | cerebellar hemisphere |
| ENSG00000109919 | MTCH2 | -5.399 | -0.036 | 6.71E-08 | 0.101 | 0.226 | 1.36E-07 | 12 | 24 | 24 | nucleus accumbens |
| ENSG00000109919 | MTCH2 | -5.388 | -0.240 | 7.13E-08 | 0.002 | 0.118 | 2.91E-05 | 9 | 18 | 18 | caudate |
| ENSG00000109919 | MTCH2 | -5.376 | -0.038 | 7.61E-08 | 0.080 | 0.121 | 4.08E-04 | 18 | 22 | 22 | anterior cingulate cortex |
| ENSG00000165915 | SLC39A13 | -5.355 | -0.071 | 8.54E-08 | 0.022 | 0.039 | 3.35E-02 | 4 | 5 | 5 | caudate |
| ENSG00000109919 | MTCH2 | -5.298 | -0.031 | 1.17E-07 | 0.121 | 0.280 | 4.48E-10 | 33 | 54 | 54 | cortex |
| ENSG00000165916 | PSMC3 | -5.229 | -0.036 | 1.71E-07 | 0.093 | 0.078 | 3.67E-03 | 38 | 63 | 63 | cerebellar hemisphere |
| ENSG00000165915 | SLC39A13 | -5.143 | -0.040 | 2.70E-07 | 0.071 | 0.064 | 4.10E-03 | 22 | 39 | 39 | cerebellum |
| ENSG00000233276 | GPX1 | 5.094 | 0.030 | 3.51E-07 | 0.148 | 0.215 | 1.45E-07 | 49 | 55 | 55 | prefrontal cortex |
| ENSG00000165915 | SLC39A13 | -5.037 | -0.068 | 4.74E-07 | 0.021 | 0.052 | 2.66E-02 | 11 | 18 | 18 | anterior cingulate cortex |
| ENSG00000172247 | C1QTNF4 | 4.943 | 0.048 | 7.70E-07 | 0.050 | 0.111 | 5.64E-04 | 30 | 46 | 46 | cerebellar hemisphere |
| ENSG00000233276 | GPX1 | 4.937 | 0.041 | 7.92E-07 | 0.072 | 0.183 | 1.46E-05 | 66 | 74 | 74 | anterior cingulate cortex |
| ENSG00000115947 | ORC4 | 4.915 | 0.049 | 8.88E-07 | 0.050 | 0.131 | 1.57E-04 | 14 | 20 | 20 | cerebellar hemisphere |
| ENSG00000213619 | NDUFS3 | -4.849 | -0.033 | 1.24E-06 | 0.094 | 0.121 | 1.35E-04 | 25 | 51 | 51 | cerebellar hemisphere |

Gene name = as listed by the Transcriptome Model, generally extracted from Genquant (http://www.gencodegenes.org/); Z score = summary PrediXcan's association

**Supplementary Table 5.** Characteristics of genotyped BioVU patients with lipid measurements

|  | HDL | LDL | TG |
| --- | --- | --- | --- |
| Number of individuals | 10722 | 10492 | 11012 |
| Number of observations | 74171 | 68737 | 77812 |
| Median lab value, mean (sd) | 49.1 (17.3) | 99.7 (32.6) | 149.7 (92.3) |
| Age in years at median lab value, mean (sd) | 59.4 (13.8) | 59.6 (13.8) | 59.1 (13.9) |
| Multiple observations |  |  |  |
| Number of individuals with >1 observation (%) | 8160 (76.1) | 7918 (75.5) | 8366 (76.0) |
| Number of observations per individual, mean (sd) | 8.8 (8.5) | 8.4 (8.1) | 9.0 (9.4) |
| Number of years between first and last observations, mean (sd) | 7.7 (5.4) | 7.3 (5.0) | 7.6 (5.4) |
| Lab value range within an individual, mean (sd) | 20.1 (14.4) | 57.3 (39.6) | 147.2 (145.0) |
| Median absolute deviation within an individual, mean (sd) | 6.5 (4.9) | 18.4 (13.5) | 44.5 (44.2) |
| Anti-lipemic medications |  |  |  |
| Number of individuals with pre-medication lab values (%) | 6742 (62.9) | 6455 (61.5) | 7060 (64.1) |
| Number of pre-medication observations ( % of all observations) | 23686 (31.9) | 21434 (31.2) | 26939 (34.6) |
| Median of pre-medication lab value, mean (sd) | 52.0 (18.3) | 115.2 (35.2) | 153.0 (101.5) |

**Supplementary Table 6.**
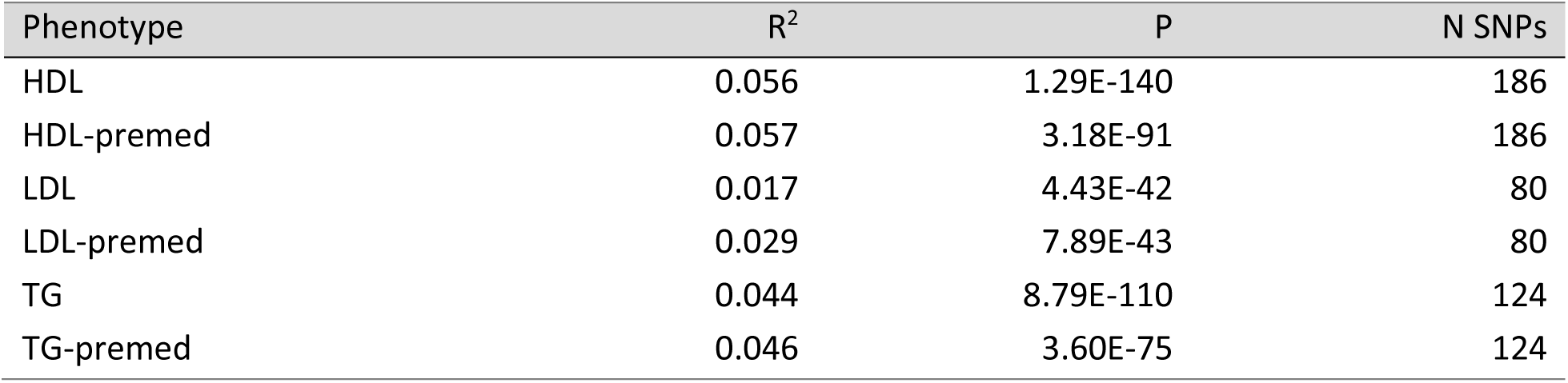
Validation of EHR-derived lipid values. Polygenic scores for HDL, LDL, and TG constructed from SNPs below pT=5×10-8 in the discovery sample were associated with the same trait in BioVU.

**Supplementary Table 7.** Prediction of lipid levels in BioVU using polygenic scores for CAD and loneliness. We identified the best fit p-value threshold for each trait pair by iterating over thresholds from 5 × 10-8 to 0.5 in increments of 5 × 10-4. R2 is the proportion of variance in the trait explained by the polygenic score, p-value is its strength of association, and N SNPs denotes the number of SNPs included in the polygenic score at a given p-value threshold.

| Base | Phenotype | Threshold | $R^2$ | $p$-value | N SNPs |
| --- | --- | --- | --- | --- | --- |
| CAD | HDL | 1 | 0.00343 | 6.96E-10 | 121732 |
| CAD | LDL-premed | 5.00E-08 | 0.00277 | 2.30E-05 | 42 |
| CAD | TG | 0.0169501 | 0.00220 | 7.52E-07 | 7636 |
| CAD | Height | 0.00085005 | 0.00052 | 2.40E-03 | 931 |
| loneliness | HDL | 0.16985 | 0.00157 | 2.99E-05 | 35861 |
| loneliness | LDL-premed | 0.00055005 | 0.00016 | 3.13E-01 | 832 |
| loneliness | TG | 0.48185 | 0.00161 | 2.40E-05 | 68271 |
| loneliness | Height | 0.0569001 | 0.00017 | 8.69E-02 | 17438 |

**Supplementary Table 10:** Cochran’s heterogeneity statistic for Inverse Variance Weighted (IVW) bidirectional two-sample Mendelian randomization analyses

| Exposure | Outcome | *n* | Cochran's Q |  |
| --- | --- | --- | --- | --- |
|  |  | SNPs | Q | *p* |
| Loneliness | BMI | 9 | 60.515 | 3.69E-10 |
| Loneliness | Body fat | 9 | 25.294 | 0.001 |
| Loneliness | Triglycerides | 9 | 27.321 | 0.001 |
| Loneliness | CAD | 12 | 16.231 | 0.133 |
| BMI | Loneliness | 53 | 120.142 | 2.59E-07 |
| Body fat | Loneliness | 10 | 14.308 | 0.112 |
| Triglycerides | Loneliness | 41 | 46.209 | 0.231 |
| CAD | Loneliness | 26 | 32.141 | 0.227 |

**Supplementary Table 11:** I-squared statistic

| Exposure | Outcome | *n* | I<sup>2</sup> |
| --- | --- | --- | --- |
|  |  | SNPs |  |
| Loneliness | BMI | 9 | 0.59 |
| Loneliness | Body fat | 9 | 0.88 |
| Loneliness | Triglycerides | 9 | 0.00 |
| Loneliness | CAD | 12 | 0.00 |
| BMI | Loneliness | 53 | 0.91 |
| Body fat | Loneliness | 10 | 0.60 |
| Triglycerides | Loneliness | 41 | 0.98 |
| CAD | Loneliness | 26 | 0.86 |

I<sup>2</sup> quantifies heterogeneity between the genetic variants in an instrument and indicates whether the 'NO Measurement Error' (NOME) assumption has been violated. If I<sup>2</sup> is smaller than 0.9, the NOME assumption is likely to be violated.

**Supplementary Table 12:** MR-Egger intercept, indicating horizontal pleiotropy, for bidirectional two-sample Mendelian randomization analyses

| Exposure | Outcome | *n*<br>SNPs | Egger intercept |  |  |
| --- | --- | --- | --- | --- | --- |
|  |  |  | intercept | *SE* | *p* |
| Loneliness | BMI | 9 | -0.020* | 0.033 | 0.556 |
| Loneliness | Body fat | 9 | -0.010* | 0.034 | 0.771 |
| Loneliness | Triglycerides | 9 | *n.a.* | *n.a.* | *n.a.* |
| Loneliness | CAD | 12 | *n.a.* | *n.a.* | *n.a.* |
| BMI | Loneliness | 53 | 0.000 | 0.002 | 0.940 |
| Body fat | Loneliness | 10 | -0.002* | 0.009 | 0.797 |
| Triglycerides | Loneliness | 41 | 0.000 | 0.001 | 0.817 |
| CAD | Loneliness | 26 | 0.003* | 0.002 | 0.116 |

\*Egger-SIMEX correction applied because of low $I^2$ estimates. *n.a.* = $I^2$ estimates too low to give reliable results for MR-Egger.

**Supplementary Figure 1:**
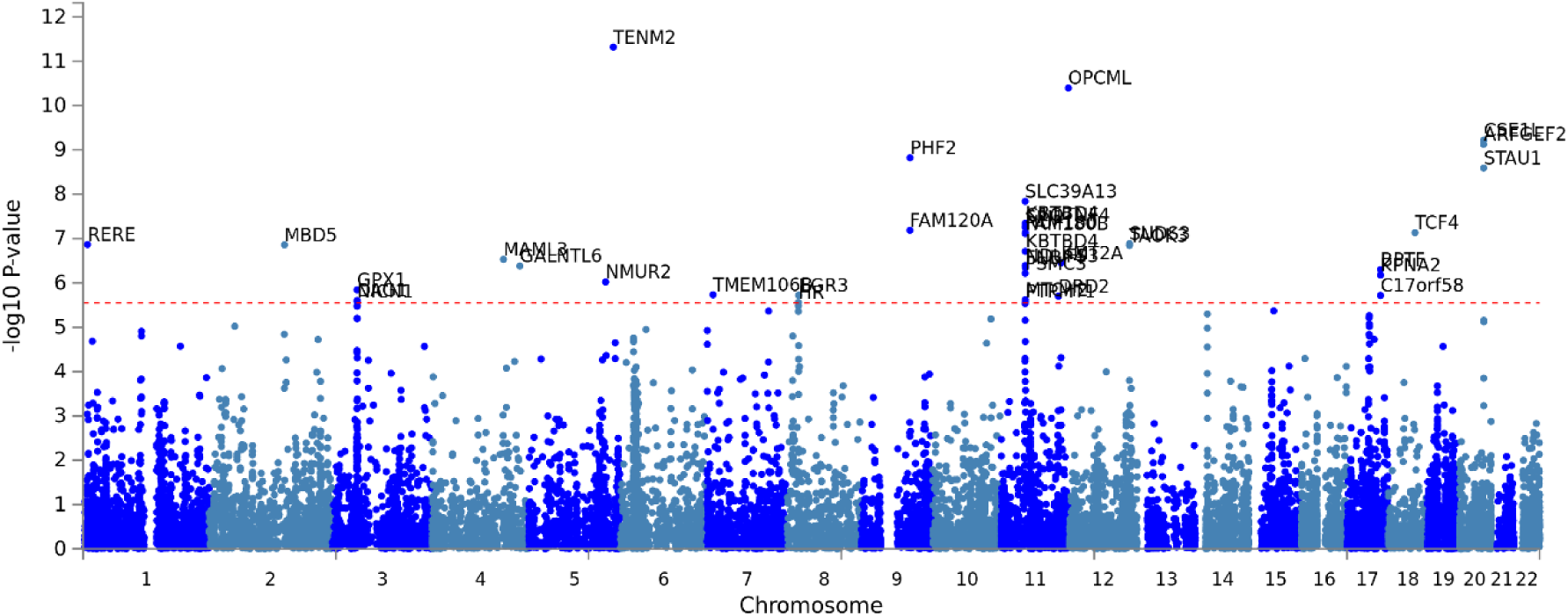
Manhattan plot of the gene-based analysis showing 38 significantly associated genes

**Supplementary Figure 2.**
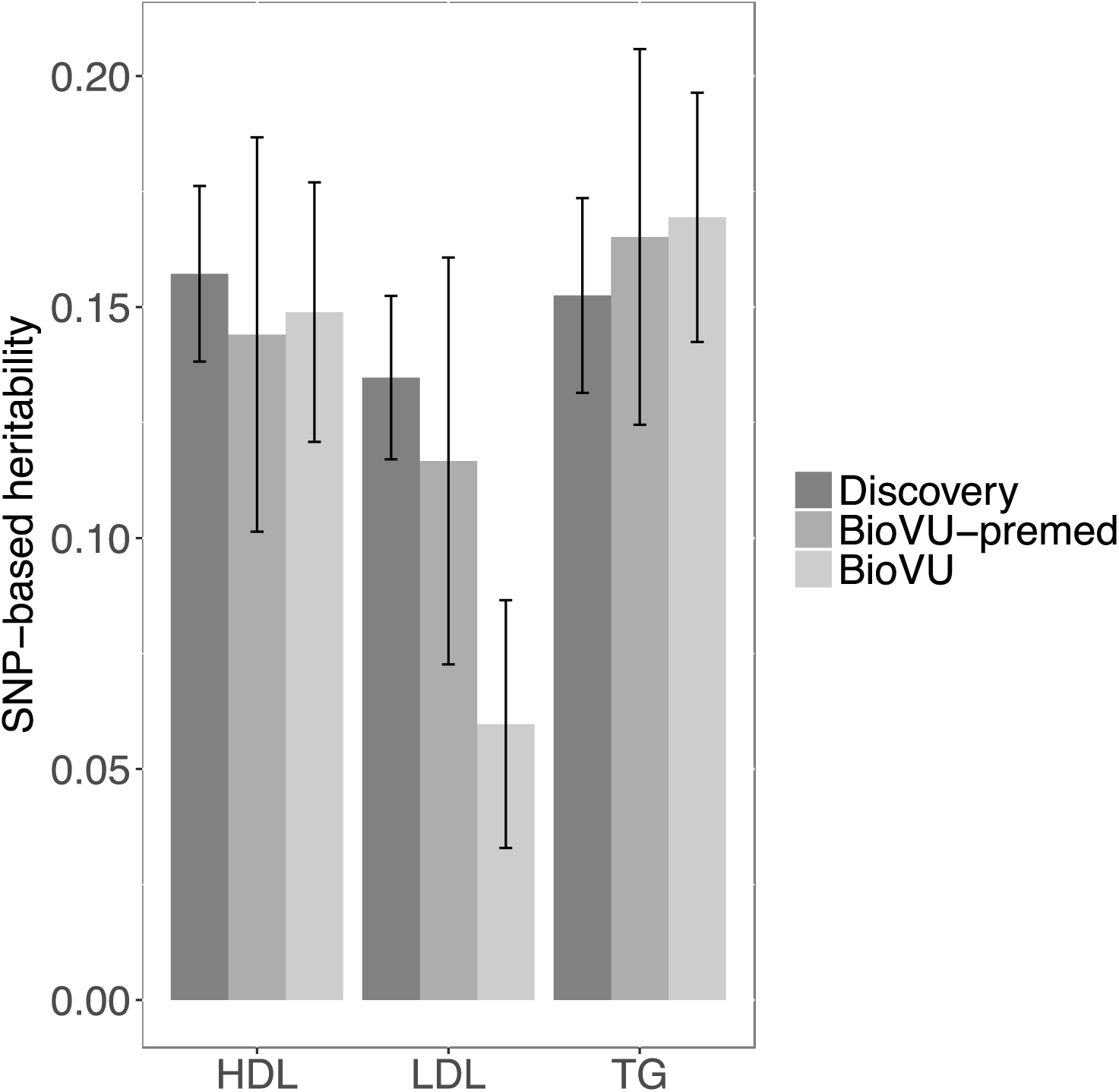
SNP-based heritability (h^2^_SNP_) of lipid values in BioVU and in the discovery dataset (Global Lipid Genetics Consortium). BioVU h^2^_SNP_ values were estimated by restricted maximum likelihood models in GCTA while discovery dataset h^2^_SNP_ values were estimated by LD score regression and extracted from LD Hub (http://ldsc.broadinstitute.org/lookup/).

**Supplementary Figure 3.**
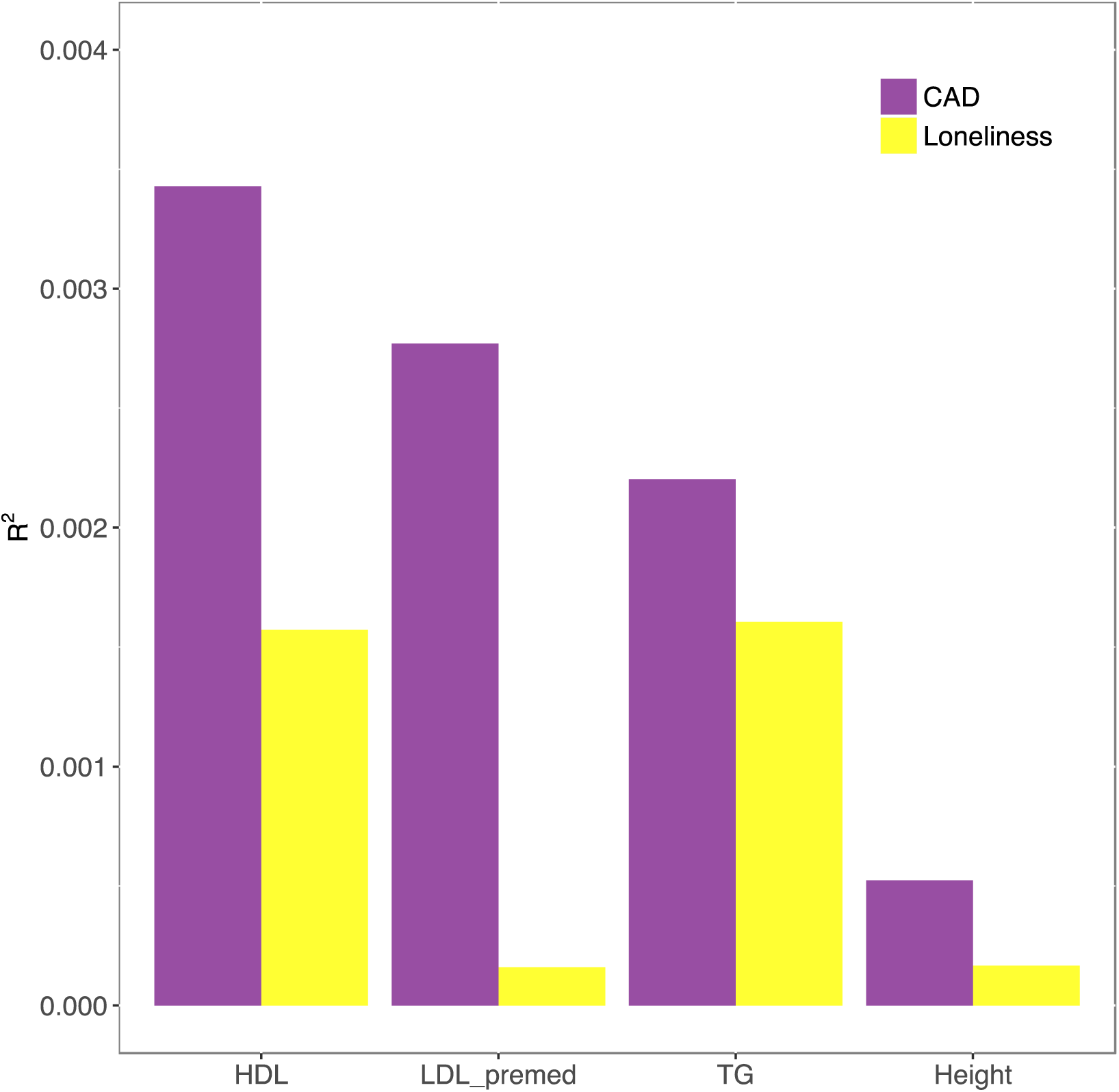
Polygenic scores for loneliness are associated with lipid levels in BioVU. The proportion of variability explained (R2) by the loneliness polygenic scores is similar to that of a CAD polygenic scores, while neither polygenic scores are associated with height (negative control).

**Supplementary Figure 4.**
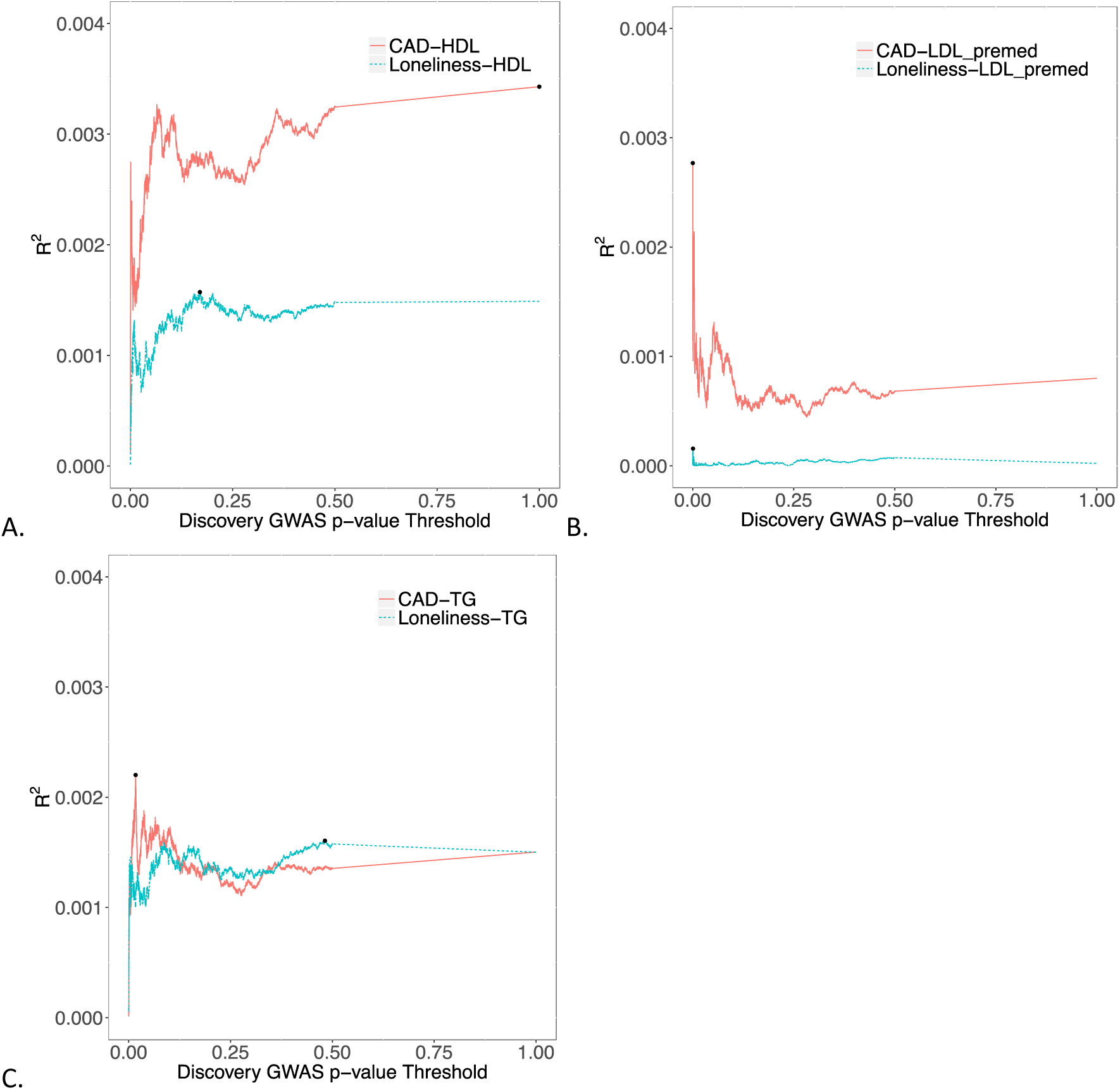
High-resolution scoring of polygenic scores for CAD and loneliness into HDL (A), pre-medication LDL values (B), and triglycerides (C). We identified the best-fit pT for each trait pair (black dot) by iterating over p-value thresholds from 5 × 10-8 to 0.5 in increments of 5 × 10-4.

**Supplementary Figure 5:**
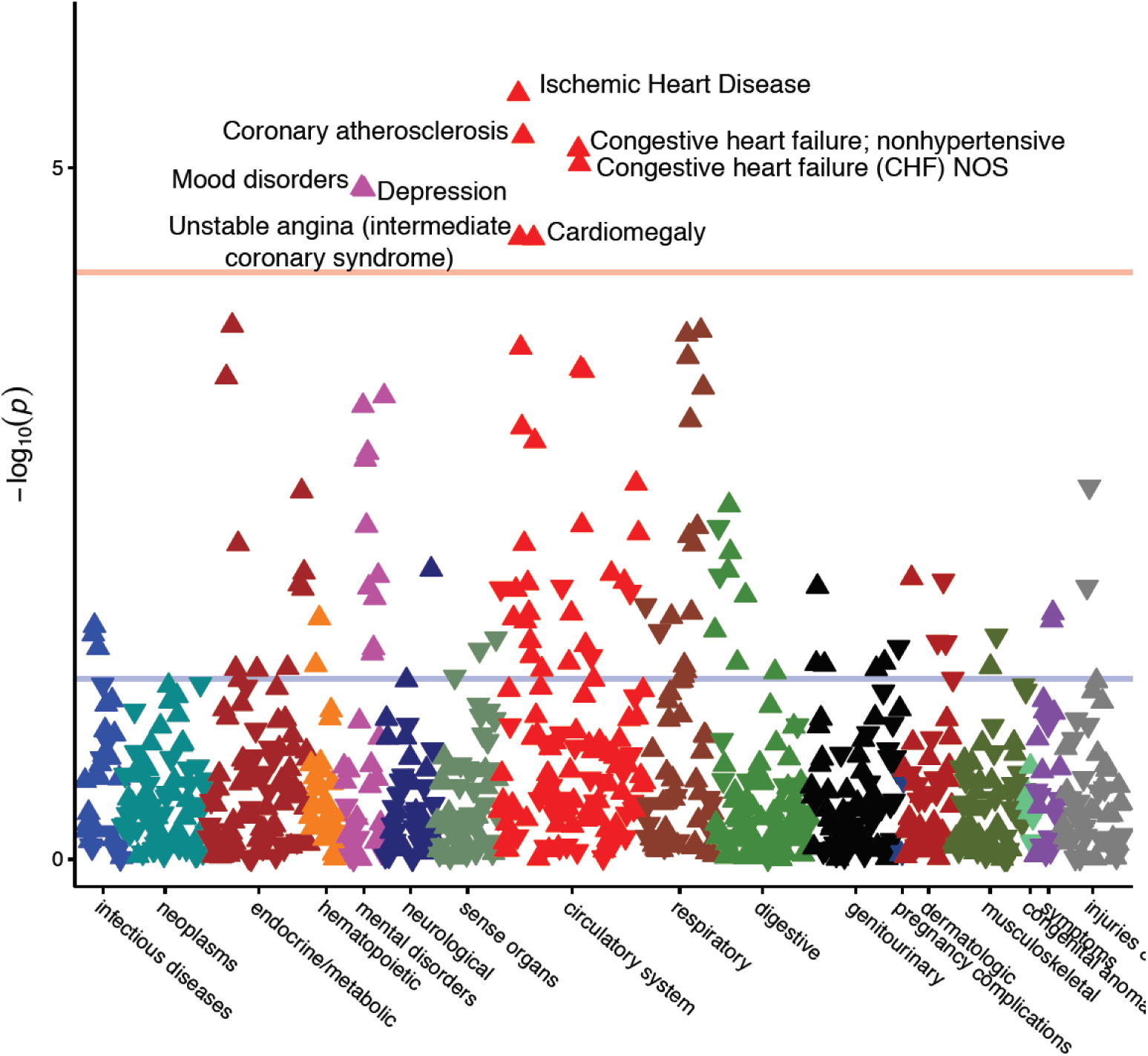
Results of the Phewas on the polygenic score for loneliness, corrected for gender, age, first 10 PCs, and batch. The loneliness polygenic score used here is constructed using only SNPs that reach p<.05 in the GWAS meta-analysis.

## Supplementary Figures on Mendelian Randomization results

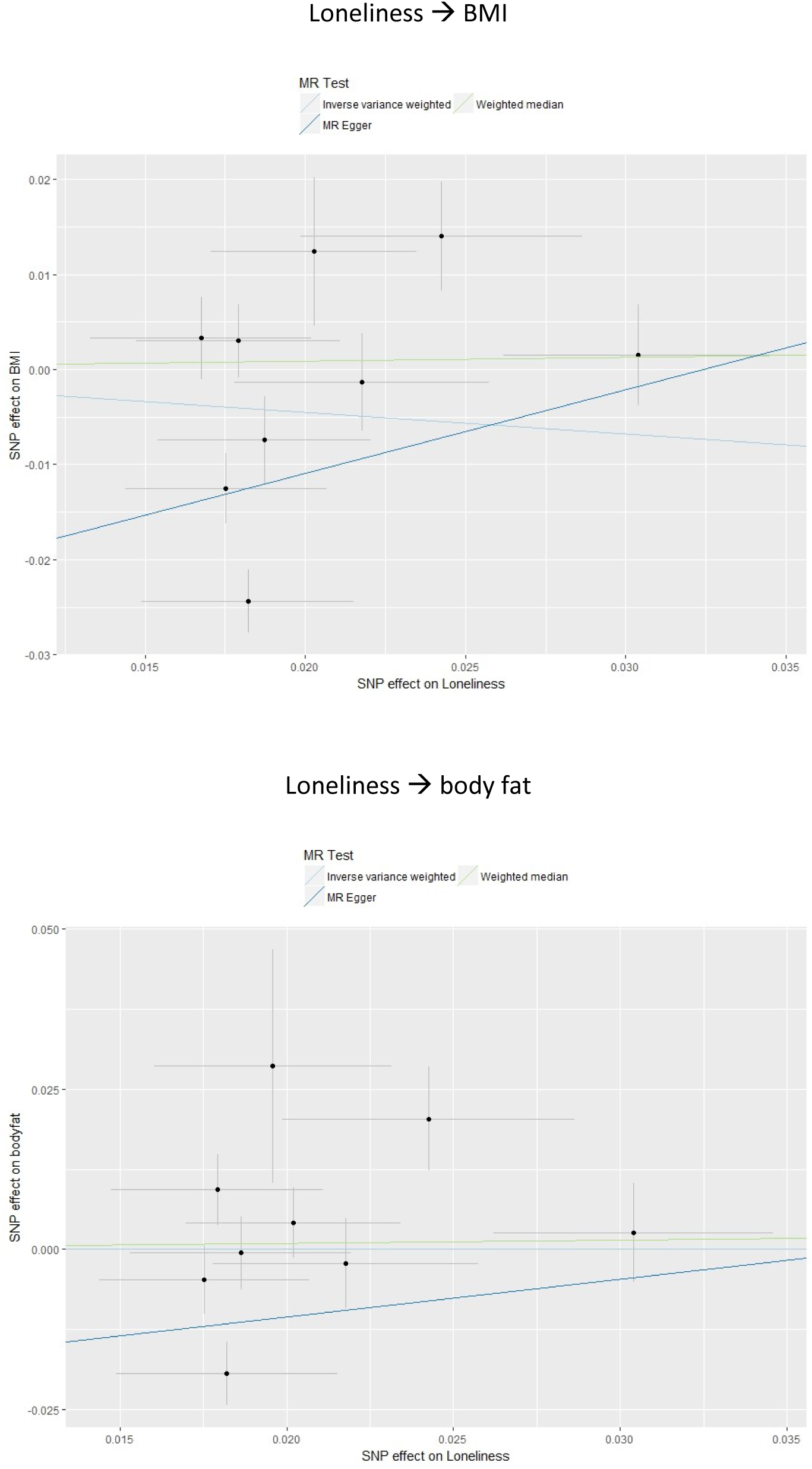
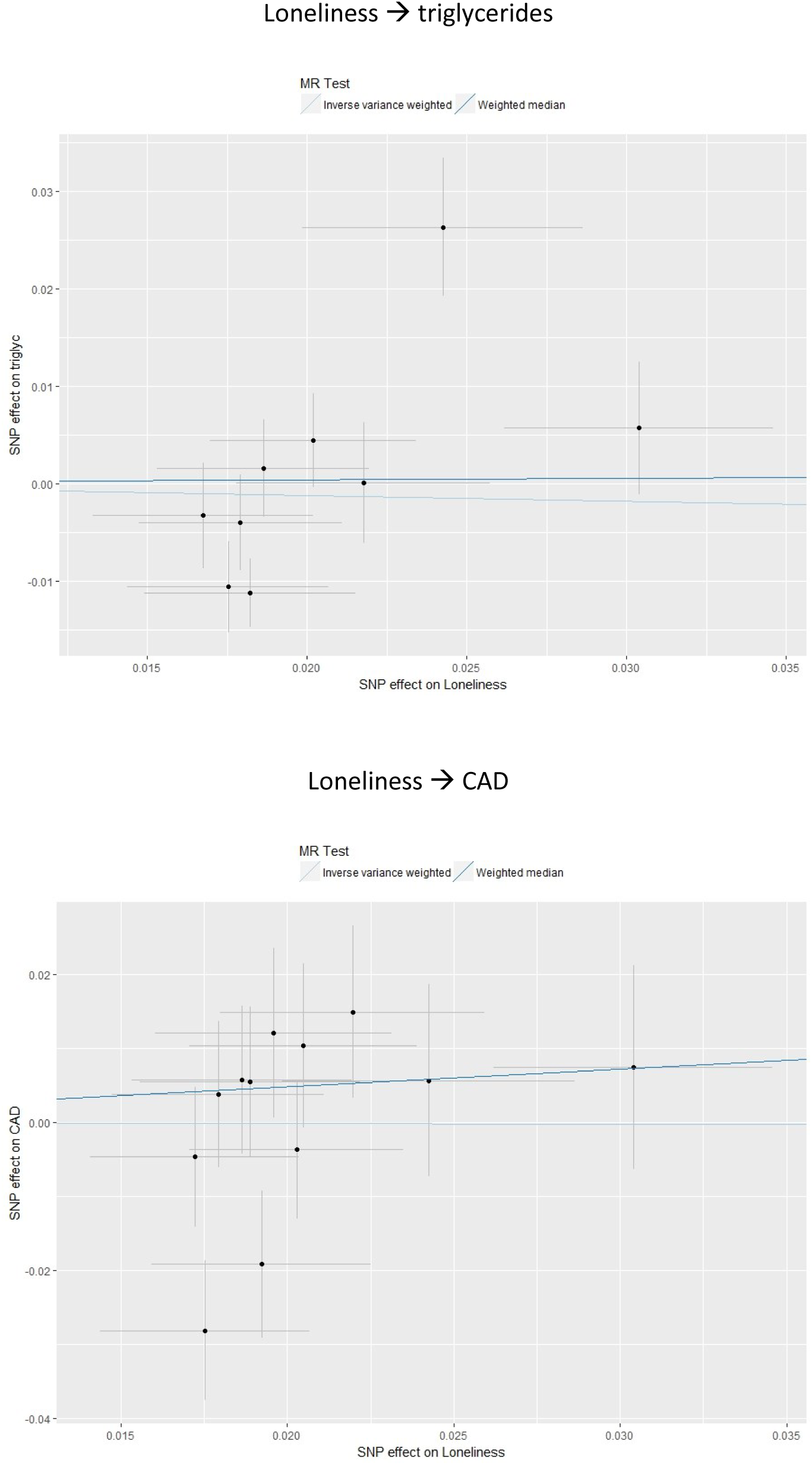
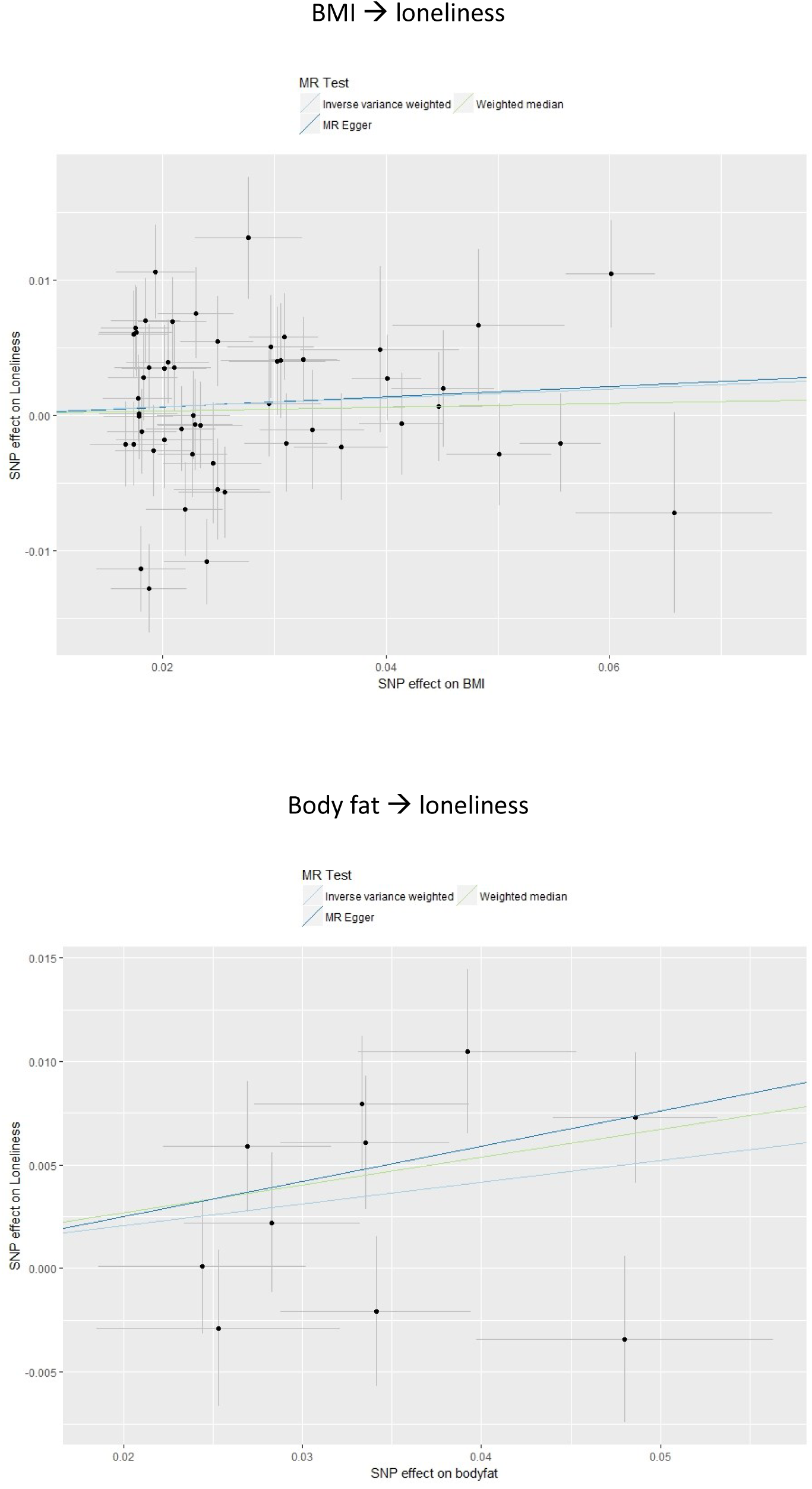
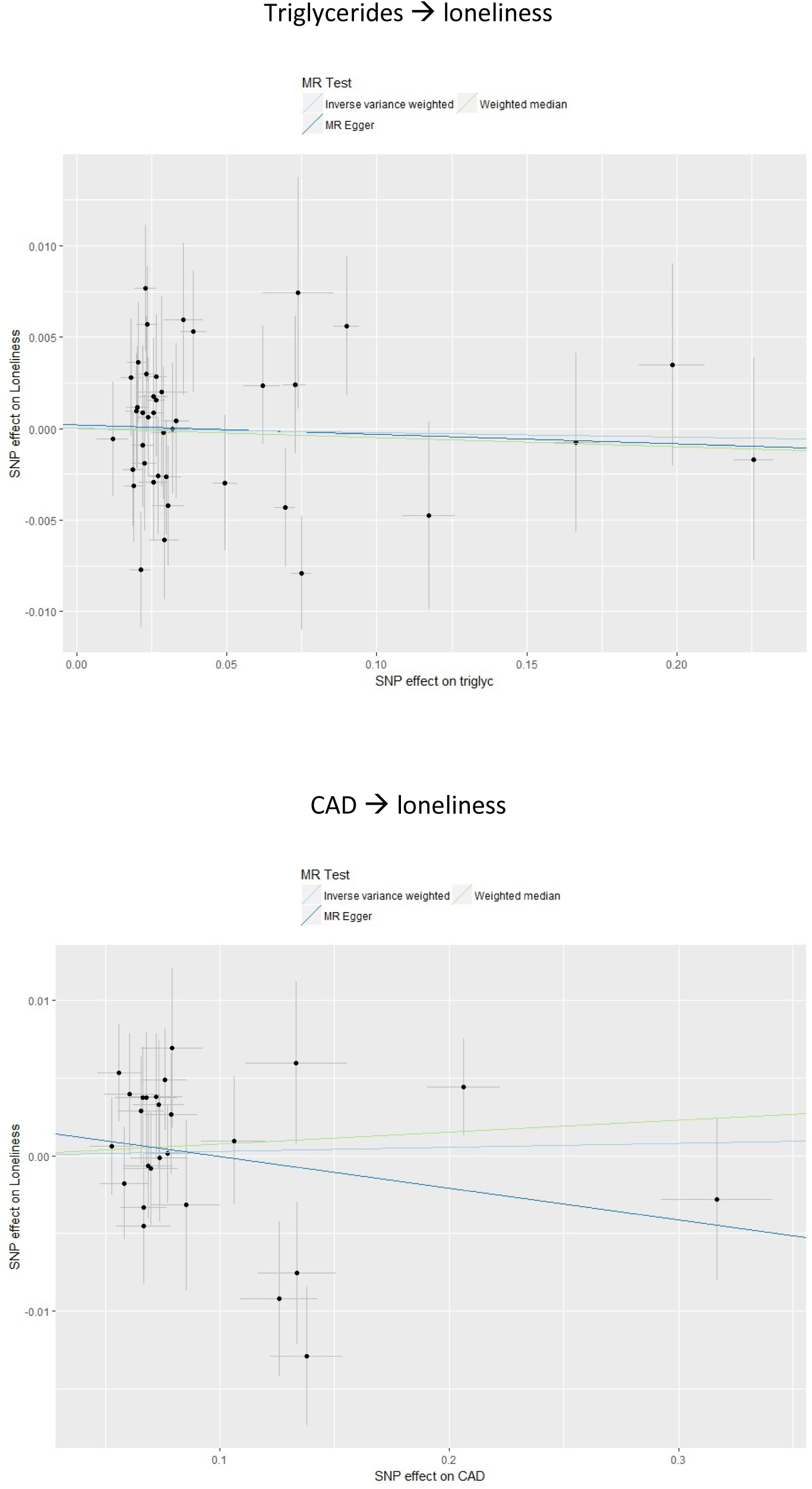

